# Geometry-driven migration efficiency of autonomous epithelial cell clusters

**DOI:** 10.1101/2022.07.17.500364

**Authors:** Eléonore Vercurysse, David B. Brückner, Manuel Gómez-González, Alexandre Remson, Marine Luciano, Yohalie Kalukula, Leone Rossetti, Xavier Trepat, Edouard Hannezo, Sylvain Gabriele

**Affiliations:** Mechanobiology & Biomaterials group, Interfaces and Complex Fluids Laboratory, Research Institute for Biosciences, CIRMAP, University of Mons, Place du Parc, 20 B-7000 Mons, Belgium; Institute for Science and Technology Austria. Am Campus 1, A-3400 Klosterneuburg, Austria; Institute for Bioengineering of Catalonia (IBEC), The Barcelona Institute for Science and Technology (BIST), 08028 Barcelona, Spain; Centro de Investigación Biomédica en Red en Bioingeniería, Biomateriales y Nanomedicina (CIBER-BBN), 08028 Barcelona, Spain; Facultat de Medicina, Universitat de Barcelona, 08036 Barcelona, Spain; Institució Catalana de Recerca i Estudis Avançats (ICREA), Barcelona, Spain

## Abstract

The directed migration of epithelial cell collectives through coordinated movements plays a crucial role in various physiological and pathological processes and is increasingly understood at the level of large confluent monolayers. However, numerous processes rely on the migration of small groups of polarized epithelial clusters in complex environments, and their responses to external geometries remain poorly understood. To address this, we cultivated primary epithelial keratocyte tissues on adhesive microstripes, creating autonomous epithelial clusters with well-defined geometries. We showed that their migration efficiency is strongly influenced by the contact geometry, and the orientation of cell-cell contacts with respect to the direction of migration. To elucidate the underlying mechanisms, we systematically explored possible cell-cell interactions using a minimal active matter model. Our investigations revealed that a combination of velocity and polarity alignment with contact regulation of locomotion captures the experimental data, which we then validated via force and intracellular stress measurements. Furthermore, we predict that this combination of rules enables efficient navigation in complex geometries, which we confirm experimentally. Altogether, our findings provide a conceptual framework for extracting interaction rules governing the behavior of active systems interacting with physical boundaries, as well as designing principles for collective navigation in complex microenvironments.

---

Collective cell migration is a fundamental process in many physiological and pathological events, such as wound healing^1^, cancer metastasis^2 3 4^ and morphogenesis^5 6 7^. Although the migration behavior of large-scale epithelial sheets has been under intense investigation in the past decades both *in vivo*^6^ and *in vitro*^8 9^, a number of physiologically relevant examples involve instead small autonomous clusters migrating with complex geometries and boundary conditions^10 11 12^. For instance, *in vivo* experiments in *Drosophila* indicated that the migration of small groups of few cells of approximately 20 µm in border cell migration of *Drosophila* to larger groups of approximately 100 µm in the posterior lateral line primordium of zebrafish^6 13^. In addition, highly polarized and persistent locomoting tumoral clusters of up to eight cells were observed in patients with epithelial-originating cancers or carcinomas^2 14^ and in mesenchymal and epithelial cancer explants originating from various organs^3 15^. To understand cell cluster migration from a biophysical perspective, *in vitro* studies of cell clusters using micropatterned confinements have focussed on cluster rotations ^16^, escape from an initial confinement ^17^, and invasion into free areas ^18 19 20 21 22 23 24^. However, the dynamics and underlying interactions that control the migration of autonomous epithelial clusters remain poorly understood, as does the key question of how cells can collectively navigate more complex geometric environments.

The migration of epithelial clusters requires front-rear polarization at the single-cell level through the formation of cryptic lamellipodia and the establishment of a large-scale coupling between cells through axial and lateral intercellular adhesive interactions^8 25 26 27 10 28^. Different interaction rules have been proposed to describe the mechanisms underlying collective migration in large epithelial monolayers, such as polarity alignment (PA), velocity alignment (VA), stress-polarity coupling (SPC) and contact regulation of locomotion (CRL)^29 30^ ^31 32 33 34 35^. From an active matter perspective, many of these possible interaction types give very similar predictions for the collective behavior of large monolayers^36^ making it hard to test each of them rigorously. Furthermore, understanding how these interactions allow cell clusters to collectively navigate complex geometrical environments could give key insights into the regulation of autonomous cell cluster migration. Studying the behavior of small clusters of cells and their response to well-defined geometrical boundary conditions could therefore give key insights in the regulation of collective motion through cell-cell interactions^27 18 20 37 38 39^. However, deciphering their individual role in the migration of small epithelial clusters remains challenging both theoretically and experimentally.

## Cluster migration depends critically on lateral, but not axial, adhesive contacts

To tackle this, we designed a set of experiments to systematically vary the size and aspect ratio of small epithelial cell clusters. We used fish epidermal keratocytes which were directly derived from primary explant tissue to study the collective dynamics of highly polarized cell clusters. Keratocytes are considered as an excellent model for cell migration, as they are characterized by fast and persistent locomotion, a simple and stable shape and strong cell-cell interactions. To generate cell trains, fish scales containing primary epithelial monolayers were placed on flat substrates functionalized with adhesive microstripes of fibronectin of various widths ^19^ (**Fig. 1a**). While such monolayers remained cohesive on homogeneous surfaces, the presence of microstripes lead to the fragmentation of the tissue into cell trains. Specifically, after few hours of explant culture, a large monolayer of primary epithelial keratocytes grew on adhesive micropatterns and formed fingers of cells that broke down into autonomous epithelial clusters of varying dimensions. To better understand the fragmentation process, we quantified the stretching of individual cells within tissue fingers and the extension speed on a 15 µm microstripe (**Extended Data Fig. 1a**) upon the fragmentation (**Extended Data Fig. 1b-c**). Our results showed an increase of the intercellular distance during the fingering process, demonstrating that cell-cell adhesive bonds are extensively stretched (up to 2-3 fold), leading to local fragmentation in an autonomous cell train (**Extended Data Fig. 1d**).

**Figure 1.**
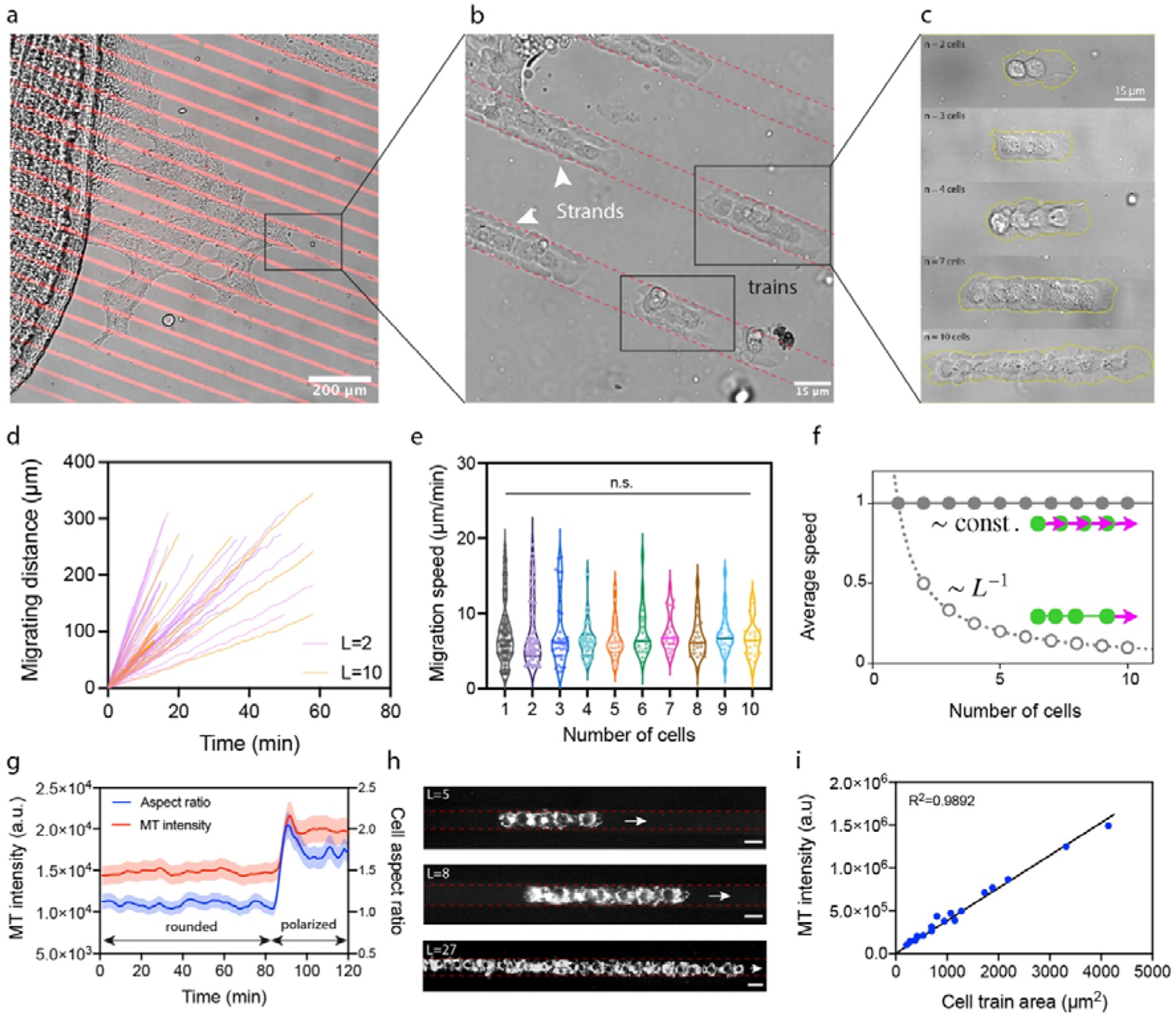
The migration velocity of cell trains is not affected by their length. **(a)** A fish scale containing epithelial keratocytes on its internal side is deposited on a PDMS substrate covered with fibronectin (FN) microstripes of 15 µm wide (in red). **(b)** The growth of the primary epithelial monolayer onto the FN microstripes leads to the formation of “strands” that break in **(c)** random-sized one-dimensional epithelial clusters called cell trains. **(d)** Representative curves of the migrating distance versus time for cell trains of L=2 (in purple) and L=10 (in orange). **(e)** Migration speed of single cells of L=1 (n=139, in black) and cell trains composed of different numbers of cells (2 ≤ L ≤ 10). Each point represents the mean migration speed of one cell train whose length (L) ranges from 2 to 10 cells, with L=2 (n=65, in purple), L=3 (n=35, in blue), L=4 (n=39, in light green), L=5 (n=23, in orange), L=6 (n=24, in green), L=7 (n=24, in pink), L=8 (n=25, in brown), L=9 (n=27, in light blue) and L=10 (n=22, in yellow). A minimum of N=5 replicates was used for each condition and the total number of cell trains (2 ≤ L ≤ 10) is 284. **(f)** Prediction of the speed as function of the train length using a model of elastically coupled active particles with either all cells polarized (solid grey dots), or with the leader cell polarized only (open dots). **(g)** Temporal evolution of MitoTracker (MT) intensity (in red) and cell aspect ratio (in blue) for individual keratocytes which underwent a transition of polarization at t=88 min and started to migrate. Data are Mean ± S.D. (n=6 with N=3 replicates). **(h)** Live mitochondria staining with MT in cell trains of different lengths (L=5, 8 and 27 cells). Scale bars in are 15 µm. **(i)** The total MT intensity in cell trains is linearly proportional their area, regardless their length. n.s. is not significant (*P*=0.8939>0.05, Kruskal-Wallis test).

We first concentrated on 15 µm wide microstripes, as it allows to form one-dimensional epithelial trains that lack lateral intercellular interactions (**Fig. 1b**). We exploited the natural variability of cell train lengths to study whether the length of these minimal epithelial clusters can affect their migration speed (**Fig. 1c**). Importantly, we found that the initial breakage from the monolayer into cell trains generates trains of different lengths, and that these trains only rarely break apart at later stages (**Supplementary Movies S1 and S2**). As reported for individual keratocytes^40^, cell trains were highly polarized and persistent (**Supplementary** Fig. 1, **Movies S3 and S4**). Interestingly, cell trains were highly compacted (**Extended Data Fig. 2a**) and their projected areas were constant over time regardless their length, demonstrating that they moved as a cohesive and single unit (**Extended Data Fig. 2b**). We found that keratocytes extended cryptic lamellipodia underneath the cell body of the preceding cell ^41^ (**Extended Data Fig. 3** and **Supplementary Movies S5**), which have been shown to be involved in polarization and cell sheet movement^42^, and can explain the compacted morphology of cell trains. We then tracked the migration of cell trains of different lengths (2≤L≤10 cells) to determine whether their migration speed was modulated by the number (L) of constitutive cells (**Fig. 1d**). Our findings showed that the migration speed of cell trains of different lengths (2≤L≤10) was similar to the speed of individual cells on 15 µm wide microstripes^40^ and did not depend on the number of cells that composed the train (**Fig. 1e** and **Supplementary** Fig. 2), even for longer cell trains with 11≤L≤18 (**Extended Data Figs. 4a**). This suggests that intercellular axial contacts – defined as front to rear cell contacts in the direction of migration – do not influence global migration efficiency, which is in stark contrast to the much lower cell velocities of tissue-level cell migration (characterized by both axial and lateral contacts, **Extended Data Figs. 4b-d**). These findings also contrast with the classic view that cells far from the leading edge would behave as less migratory “follower” cells, as this would theoretically predict a decrease of global velocity with the train length L (with a 1/L trend, see **Fig. 1f**), instead suggesting a model where all cells are equally polarized (**Fig. 1f**). To confirm this, we used both the mitochondrial potential membrane and the localization of the Golgi complex as a readout of the cell polarization. Indeed, it was shown that cell polarization requires increased cellular energy^43^ and that the mitochondrial activity promotes cell migration by supporting membrane protrusion and modulating F-actin polymerization^44^. Stationary and unpolarized keratocytes exhibited a rounded shape and a low MitoTracker (MT) intensity that increased immediately with their polarization (**Fig. 1g**, **Extended Data Fig. 5a-b** and **Supplementary Movie S6**). Interestingly, all cells within migrating cell trains of different lengths showed a large MT signal (**Fig. 1h**), reflecting a high level of mitochondrial activity that was constant over time during migration of individual cells (**Extended Data Fig. 5c**) and cell trains (**Extended Data Fig. 5d-e** and **Supplementary Movie S7**). In addition, our results indicated a linear correlation between MT intensity and cell train areas (Fig. 1h-i and **Extended Data Fig. 5f**). In contrast with individual keratocytes where microtubules are densely packed in the cell body and wrapped around the nucleus by forming a cage with no preferential organization ^40 45^, we found that microtubules in cell trains were mostly aligned with the axis of migration and started to extend in the cryptic lamellipodia (**Extended Data Fig. 3**), suggesting a polarized state ^46^. Furthermore, in agreement with previous observations on polarized individual cells confined on narrow microstripes ^47^, the Golgi complex of individual cells in cell trains was positioned behind the nucleus and at few microns away from it (**Extended Data Fig. 6**), confirming the polarization of all cells within a cell train. Altogether, these findings indicate that all members of cell trains are metabolically active, polarized and contribute to the migration process, driving length-independent collective motion of cell trains.

We next sought to investigate the role of lateral adhesive interactions in collective migration, which were missing in one-dimensional cell trains. For this, we examined larger clusters of primary epithelial cells that formed on 30, 45 and 100 µm wide microstripes from an explant (**Fig. 2a**). Epithelial monolayers grew up on FN microstripes and fragmented in epithelial clusters of controlled widths (**Fig. 2b**) that were composed of compacted cells with cryptic lamellipodia (**Extended Data Fig. 3**), as observed in the primary epithelial monolayer. Time-lapse experiments indicated that these migrating epithelial clusters of various widths were highly persistent (**Supplementary Movies S8 and S9**). The constituent cells of these clusters exhibited directed motion along the main axis of the microstripes (**Fig. 2c**), suggesting that the relative cell position within the cluster was maintained during the whole migration process, regardless the cluster width. Strikingly, widening the cell cluster to 30 µm, 45 µm and 100 µm strongly decreased the migrating velocity from 8.1±3.9 µm/min (n=59) for cell trains on 15 µm to 3.3±1.2 µm/min (n=51) on 30 µm, 2.5±0.8 µm/min (n=118) on 45 µm and 2.2±0.8 µm/min (n=64) on 100 µm (Fig. 2d), slowly converging towards the migration speed of large-scale epithelial monolayers (1.4±0.4 µm/min, **Extended Data Fig. 5f**). This was unlikely to be due to neighbor-driven confinement as we found similar cellular densities within the different epithelial cluster sizes (**Fig. 2e**). Altogether, these findings suggests that lateral cell-cell adhesive interactions have drastically different consequences – compared to axial interactions – in collective cell migration.

**Figure 2.**
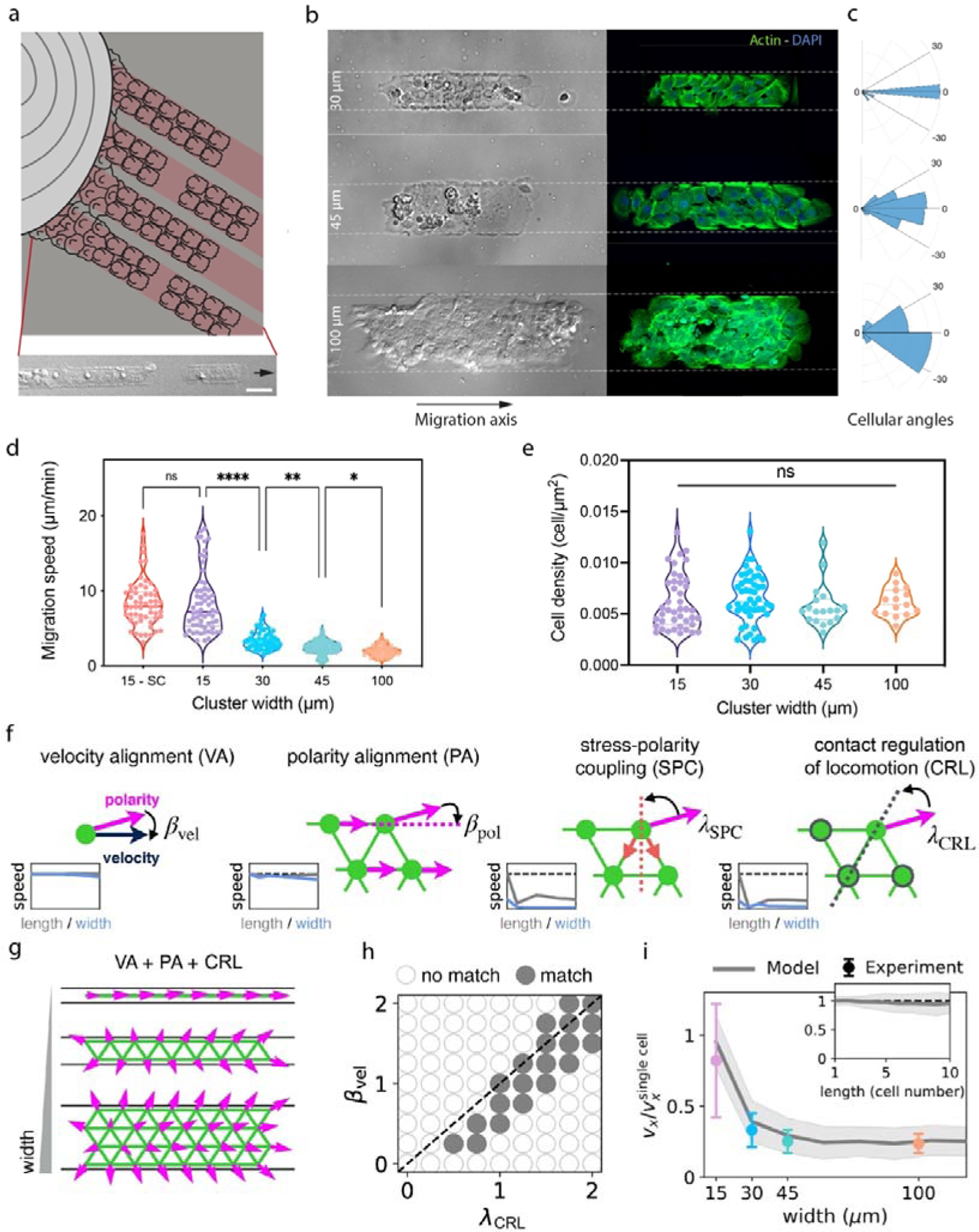
The migration velocity decreases with the epithelial cluster widening. **(a)** Sketch of the formation of autonomous epithelial clusters that detach from fingers of a primary epithelial tissue formed during its expansion on adhesive microstripes (in light red). The insert shows a representative DIC image of an epithelial cluster detached from a tissue extension on a 30 µm wide microstripe. Widening the microstripes (30, 45 and 100 µm) leads to a gradual increase of the number of cell-cell interactions. **(b)** Typical micrographs in DIC and epifluorescence mode (actin in green and DAPI in blue) of epithelial clusters formed on FN microstripes of 30, 45 and 100 µm. **(c)** Distribution of the migration angle of each individual cell within an epithelial cluster migrating during 60 min on a 30 µm (top), 45 µm (middle) and 100 µm (bottom) wide microstripe. The axis of the microstripe (horizontal axis) corresponds to 0°. **(d)** Distribution of the migration speed for single epithelial cells on 15 µm wide microstripes (SC, in pink) and epithelial clusters on 15 µm (in purple), 30 µm (in blue), 45 µm (in green) and 100 µm (in orange). **(e)** Cell density of epithelial clusters of different widths. A minimum of N=5 replicates was used for each condition. **(f)** Schematic illustration of each of the four considered cell-cell interaction mechanisms, from left to right: velocity alignment (VA), polarity alignment (PA), stress-polarity coupling (SPC), and contact regulation of locomotion (CRL), quantified by amplitudes β_vel_, β_pol_, β_SPC_, β_CRL_, respectively (**Supplementary Theory Note**). Green dots and lines represent cells and elastic links between cells, respectively, and blue, magenta and red arrows denote cell velocity, polarity, and inter-cellular stress, respectively. Insets: predicted dependence of cluster speed as a function of length (grey) and width (red) for each of the mechanisms. Dashed line indicates the speed of a single cell. **(g)** Predicted polarity fields as a function of cluster width for the combination of VA, PA and CRL. **(h)** Phase diagram indicating where the VA+PA+CRL model captures the qualitative trends of speeds independent of length and decreasing with width (solid grey dots), as a function of the VA and CRL strength, for fixed β_pol_ = 1.5. Similar trends are observed for other β_pol_ (**Supplementary Theory Note**). **(i)** Predicted speed dependence as a function of width (red) and length (grey) for best fit parameters β_vel_ = 0.15,β_pol_ = 1.5, β_CRL_ = 0.5, compared to experiment (blue squares). Speeds in both model and experiment are normalized by the single cell speed. Error bars denote standard deviations, **p<0.01; ***p < 0.001; ****p < 0.0001 and n.s., not significant (post-hoc Dunn’s test), with p< 0.0001 for (d) and p>0.05 for (e) using Kruskal-Wallis tests.

To shed light on the role of the cell-cell adhesions in this behavior, we induced chemical disruption of cell-cell junctions with triethylene glycol diamine tetraacetic acid (EGTA) (**Extended Data Figs. 7a-b**)^48^. After EGTA treatment, we observed a fragmentation of all epithelial clusters into individual cells. Interestingly, EGTA treatments performed on 30 µm wide clusters and primary monolayers induced a speed up of the cells, whereas the cell migration speed remained constant after EGTA treatment on single cells and 15 µm wide clusters (**Extended Data Fig. 7c**). Altogether, these data argue that the establishment of an increasing number of lateral, but not axial, contacts decrease migration speed during collective migration.

## Theoretical modelling of geometry-dependent cluster migration

The observation of length-independent but width-dependent collective motion suggests an intricate interplay of cell-cell interactions with the cell cluster geometry. From a theoretical perspective, various interaction types have been proposed in the context of collective migration^36^, but it is unclear which interactions are required to capture the migration dynamics of small confined epithelial clusters. To disentangle the role of these interactions, we developed a minimal theoretical model of confined cell clusters. Based on our observations that cells in trains and clusters are polarized and exhibit few cellular re-arrangements and density variations, we modelled cells as polar particles exerting active migration forces in a direction of polarity p (defined in our system by the orientation of the lamellipodia), with each cell i connected to its neighbours *j* by elastic links (adhesions). This model is described by the non-dimensionalized equation *dx_i_/dt = ∑ F^elastic^ + p_i_(t)*. Importantly, this model allows cell velocities Importantly, this model allows cell *dx_i_/dt* and polarities *p_i_* that are not necessarily equal, due to cell-cell mechanical interactions. The evolution of the cell polarity can then be generally modelled as *dp_i_/dt = p_i_(1 - |p_i_|^2^) + F^int^+ √2nr,_i_(t)*, where the first term describes the spontaneous polarization of single cells, and r,_i_(t) is Gaussian white noise (see **Supplementary Theory Note** for details)^49^.

The general interaction term F^int^ subsumes many possible cell-cell interactions that could give rise to the intricate collective dynamics of confined cell clusters observed here (Fig. 2f). Two classes of interactions that have been particularly studied are neighbouring cells either aligning or anti-aligning relative to each other^30 36 50 51^. Indeed, in our system, two-cell collision experiments can show both alignment (**Extended Data Fig. 8a**) and anti-alignment (**Extended Data Fig. 8b**) depending on the initial configuration^52 53^. Theoretically, on the one hand, alignment is motivated by the tendency of cells to “flock” in the same direction, which can be described by i) alignment of the polarity of each individual cell to match its own velocity (velocity alignment, VA)^34 36 18^ and ii) direct polarity alignment (PA) between neighbours, in analogy to spins in magnetism^54 55^. On the other hand, a number of cell types have been shown to anti-align to counter-act the forces exerted by their neighbours, either due to i) stress-polarity coupling (SPC) where cells exert polarity forces opposite to stress applied on their cell-cell contacts^49 27 56^ or ii) contact regulation of locomotion (CRL), which controls cell polarity in response to cell-cell contacts^31 33 57^. While CRL is often considered in the setting of two isolated cells colliding and repolarizing (the typical contact inhibition of locomotion scenario ^57 58^), we consider it here in the more general case of strong adhesion between cells, which is relevant in a number of physiologically relevant situations ^58 59^. In this context, CRL lead cells to a contact-dependent polarization away from one another without necessarily breakage. Importantly, we enumerate all possible couplings allowed by symmetry within our minimal model and show that they correspond to each of these categories (**Supplementary Theory Note**). This motivates a key question: could several distinct types of interactions interplay to generate the observed nontrivial coupling to cluster geometry?

We thus proceeded to simulate clusters with varying cluster lengths and widths, under each of the possible combinations of interaction types. Generically, alignment interactions (PA or VA) lead to geometry-independent speed with all cells moving together in a specific direction, while anti-alignment interactions (SPC or CRL) induce rapidly decreasing speed with increasing length and width as clusters tend to develop a bidirectional polarity pattern (**Fig. 2g**, **Supplementary Movie S6**). However, we reasoned that combining different mechanisms – to give rise to alignment in the axial direction and anti-alignment in the lateral direction – could be a promising avenue to explain our data. Interestingly, the screen of different computational models revealed that pair-wise combinations of interactions could still not qualitatively capture our observations (**Supplementary Theory Note**, **Supplementary Movie S10**), but that the combination of VA, PA, and CRL represents the minimal interactions necessary to reproduce length-independent, but width-dependent migration (**Fig. 2h**). Qualitatively, this combination of interactions causes length-independent aligned cell motion in the axial direction, while the presence of boundaries along the lateral direction renders such an aligned state impossible, allowing lateral anti-alignment to develop, driven by the tendency from CRL of boundary cells to polarize outwards (**Fig. 2g**). Here, velocity and polarity alignment play distinct roles: while VA breaks the symmetry between axial and lateral directions, as non-zero global velocities cannot arise in the lateral direction; and PA propagates the CRL-driven laterally outwards-pointing polarity into the bulk. This causes a re-orientation of polarity at the detriment of “productive” motion which can only occur in the axial direction. In the case where VA and CRL have similar magnitudes, this interplay of geometry and interactions then gives rise to the observed trends of speed independent of length but decreasing with width (**Figs. 2h-i**, **Supplementary Movie S11**).

Beyond recapitulating the key differences between axial and lateral adhesion for the migration speed of trains, we also examine different extensions of the model, to consider the effect of adhesion strength and cell-to-cell variability on the robustness of the predictions. We used our model to check that the constant cell speed of cellular trains was not an artifact due to cells of different intrinsic speed detaching from each other. For this, we included the possibility of junctional breakage (when cell-cell distance exceeds a critical threshold *l*_c_, **SI Theory Note**, **Fig. S5**), as well as intrinsic variability in cellular migration forces (based on single-cell migration speed variability, **SI Theory Note Fig. S5**). For low adhesion, trains frequently break apart even due to intrinsic variability and noise on protrusion forces, in sharp contrast to our observations of extremely rare detachments both in trains and confluent tissues. These analyses allow us to estimate a lower bound for the adhesion force in fish keratocytes (**SI Theory Note**). Furthermore, an upper bound on adhesion forces can also be obtained from explicit modelling of the initial fingering process, where trains detach from the bulk due to the very high cellular stretches arising from the microstrip constraint (**Extended Data Fig. 1, Supplementary Movie S12** and **SI Theory Note**).

Together, these findings demonstrate how multiple types of cell-cell interactions interplay to determine the geometry-dependent migration efficiency of cell clusters. These cell clusters exhibit strong cell-cell adhesion, allowing them to remain cohesive in the presence of outward-pointing polarities.

## Testing the predicted interaction regime via traction force microscopy

Our model makes a clear and experimentally testable prediction: the outward polarization induced by CRL at the boundary and propagated by PA causes a build-up of lateral inter-cellular stress, which we predict to increase with increasing width. To test this key prediction, we performed traction force microscopy (TFM) experiments to quantify the orientation of the traction forces generated by cell clusters of different geometries. To separate axial (along the microstripe axis) and lateral (perpendicular to the microstripe axis) components of the traction stresses (**Fig. 3a**), we plotted tractions in a reference frame where the horizontal and vertical axes (*x*, *y*) correspond to length and width of the cell cluster, respectively. As shown in **Fig. 3b**, cell trains were characterized by a dipole of forces concentrated at both extremities of the cell train and directed inward toward the center of the train, implicating a strong intercellular coupling^8^. Interestingly, widening cell clusters to 45 and 100 µm led to the generation of more pronounced lateral traction forces. By calculating the strain energy as the dot product of the traction force with the displacement (**Fig. 3c**), we determined the individual contribution of axial (*E_x_*) and lateral (*E_y_*) components of the strain energy. In agreement with a dipole of forces, we found that cell trains on 15 µm wide microstripe exhibited a large axial component, which accounted for ∼96.7% of the total strain energy (**Fig. 3d**) while the lateral component was negligible (∼4.3%). Interestingly, the lateral component *E_y_* increases with width, even leading to an inversion of the major strain energy component for 100 µm wide clusters, with *E_y_*that represented ∼27.1% and ∼56.4% of the total strain energy for 45 and 100 µm wide clusters, respectively (**Fig. 3d**). As shown in **Fig. 3e**, widening cell clusters leads therefore to the inversion of the major strain energy component for 100 µm wide clusters. Taken together, these findings point toward a transition between axial and lateral contractile forces as a function of the cluster width, where the larger amount of traction forces in wider cell clusters is exerted normal to the direction of migration.

**Figure 3.**
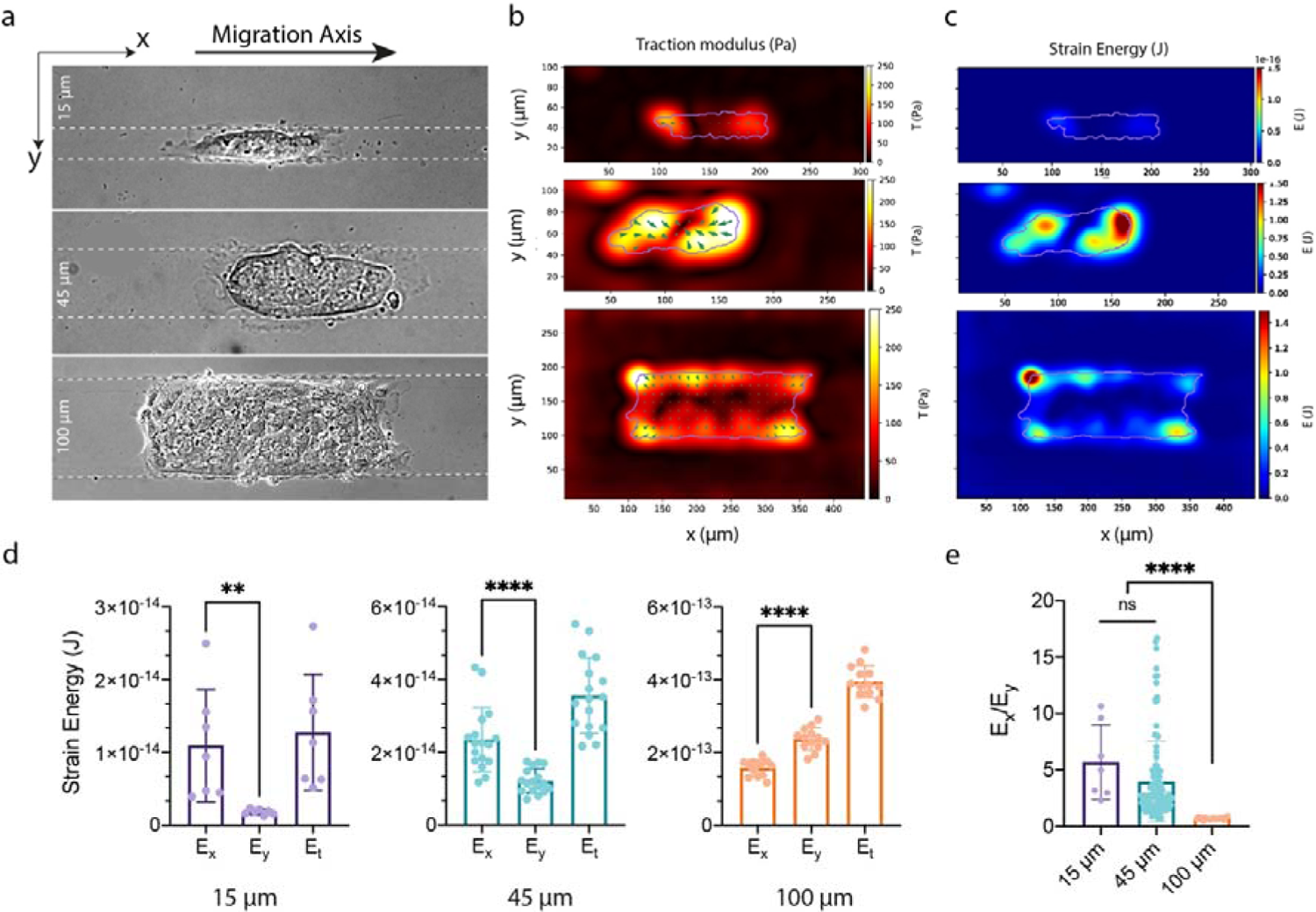
Widening epithelial clusters leads to the emergence of lateral contractile forces and lateral intracellular stresses. **(a)** Typical DIC images of epithelial clusters on 15 µm, 45 µm and 100 µm wide microstripes. **(b)** Heat map of the spatial distribution of the traction modulus and **(c)** the strain energy field exerted on the substrate during the migration of epithelial clusters of 15 µm, 45 µm and 100 µm wide. Distribution of **(d)** axial (*E_x_*) and lateral (*E_y_*) components of the total strain energy (*E_t_*) and **(e)** *E_x/_E_y_* ratio for epithelial clusters of 15 µm wide (in purple), 45 µm (in green) and 100 µm (in orange) wide. **p < 0.01, ****p < 0.0001 and n.s. not significant. Error bars denote standard deviations.

As traction forces must be balanced by internal forces transmitted within and between cells, we can infer the spatio-temporal profile of the stress tensor within the monolayer using Monolayer Stress Microscopy (MSM), and thus calculate axial (σ_xx_) and lateral (σ_yy_) components of the internal stress field^60 61^. Our model indicated a very small amount of lateral stress in cell trains of 15 µm wide, which increased with the cluster widening (**Fig. 4a**). Interestingly, we have shown by confocal microscopy that keratocyte cells extend cryptic lamellipodia against the substratum beneath cells in front of them (**Extended Data Fig. 3**). More quantitatively, our model – with parameters fully constrained based only on the speed variations as a function of cluster geometry (**Fig. 2j**) – successfully predicted a significant increase of the stress ratio σ_yy_/σ_xx_ as a function of the cluster width (**Fig. 4b**), in agreement with MSM experiments conducted on 15, 45 and 100 µm wide clusters (**Fig. 4c**). The good agreement between our theoretical (**Fig. 4b**) and experimental (**Fig. 4e**) results highlight the role of the lateral stress component in wide epithelial cluster, as well as the cooperative role of several modes of cell-cell interaction to shape the collective migration and stress profile of small cell clusters.

**Figure 4.**
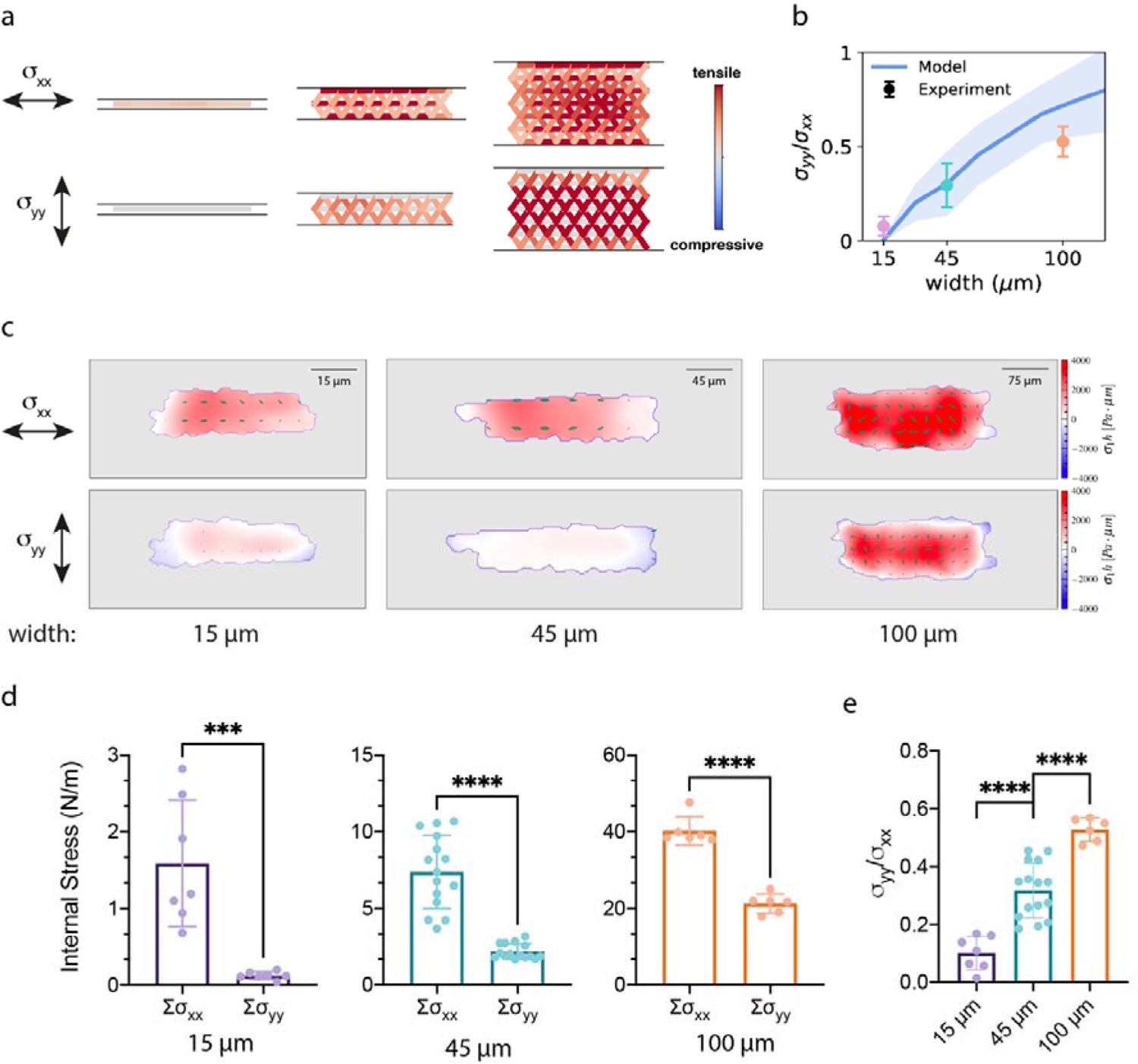
The lateral internal stress increases in wider epithelial clusters. **(a)** Theoretical predictions of axial (σ_xx_; top) and lateral (σ_yy_; bottom) components of the internal stress field for varying cluster widths (1, 3 and 7 cells, corresponding to 15, 45 and 100 µm) using the VA+PA+CRL model. **(b)**. Ratio of the lateral to axial stress σ_yy_/σ_xx_ as a function of cluster width as predicted by the model with best parameters identified based on cell speed (see Fig. 2j). **(c)** Typical maps of the spatial distribution of axial (σ_xx_) and lateral (σ_yy_) internal stress components for cell trains of 15 µm wide (in light purple) and epithelial clusters of 45 (in light green) and 100 µm (in light orange) wide. **(d)** Quantification of σ_xx_ and σ_yy_ and **(e)** σ_yy/_ σ_xx_ratio. ***p < 0.001, ****p < 0.0001 and n.s. not significant. Error bars denote standard deviations.

## Migration efficiency in response to complex boundary conditions

The cell polarization fields predicted by our model further suggests non-trivial consequences for the lamellipodial orientation, due to the competing cell-cell interactions, with both axial and lateral polarization components at the lateral boundaries. To test for the lateral components due to CRL, we imaged cell clusters that reach the end of a 1D microstrip leading to an open space (**Extended Data Fig. 9a** and **Supplementary Movie S13**). These clusters rapidly develop large lamellipodia in the lateral direction away from their neighbours, consistent with the assumption of CRL in the model and confirmed by simulations for this scenario (**Extended Data Fig. 9b** and **Supplementary Movie S14**). Importantly, even in this context, clusters do not break apart, consistent with our assumption of strong adhesions mediating mechanical and polarity interactions between cells (**Extended Data Fig. 10** and **SI Theory Note**).

This specific combination of several mechanisms with opposite effects on cell polarity raises the question of what functional consequences this could have, especially since they lead to substantial unproductive lateral stresses and thus slower collective migration. To address this question, we used our model to predict how various combinations of cell-cell interactions affect cluster behavior in response to more complex external environments. When subjected to environments with blind-ends, where cell clusters must abruptly change their polarization, we found that the repolarization behavior is strongly affected by the choice of cell-cell interactions considered in the model (**Fig. 5a-d, Extended Data Fig. 10a-f and Supplementary Movie S15**). Indeed, while VA and PA mechanisms lead to fast migration in uninterrupted straight microstripes, they result in poor repolarization abilities, while, intuitively, CRL leads to slow-moving clusters which however repolarize quickly when contact geometry changes. Interestingly, the parameter region that captured the experimental observations (intermediate values of all three interaction parameters), exhibits a seemingly optimal behavior with both fast straight migration and obstacle-driven repolarization **(Extended Data Fig. 10b**). Importantly, time-lapse experiments and quantitative cell trackings performed on epithelial clusters reaching the end of a FN stripe agree with the theoretical prediction, showing fast and global repolarization of both cell trains and clusters (**Extended Data Fig. 10d-f)**.

**Figure 5.**
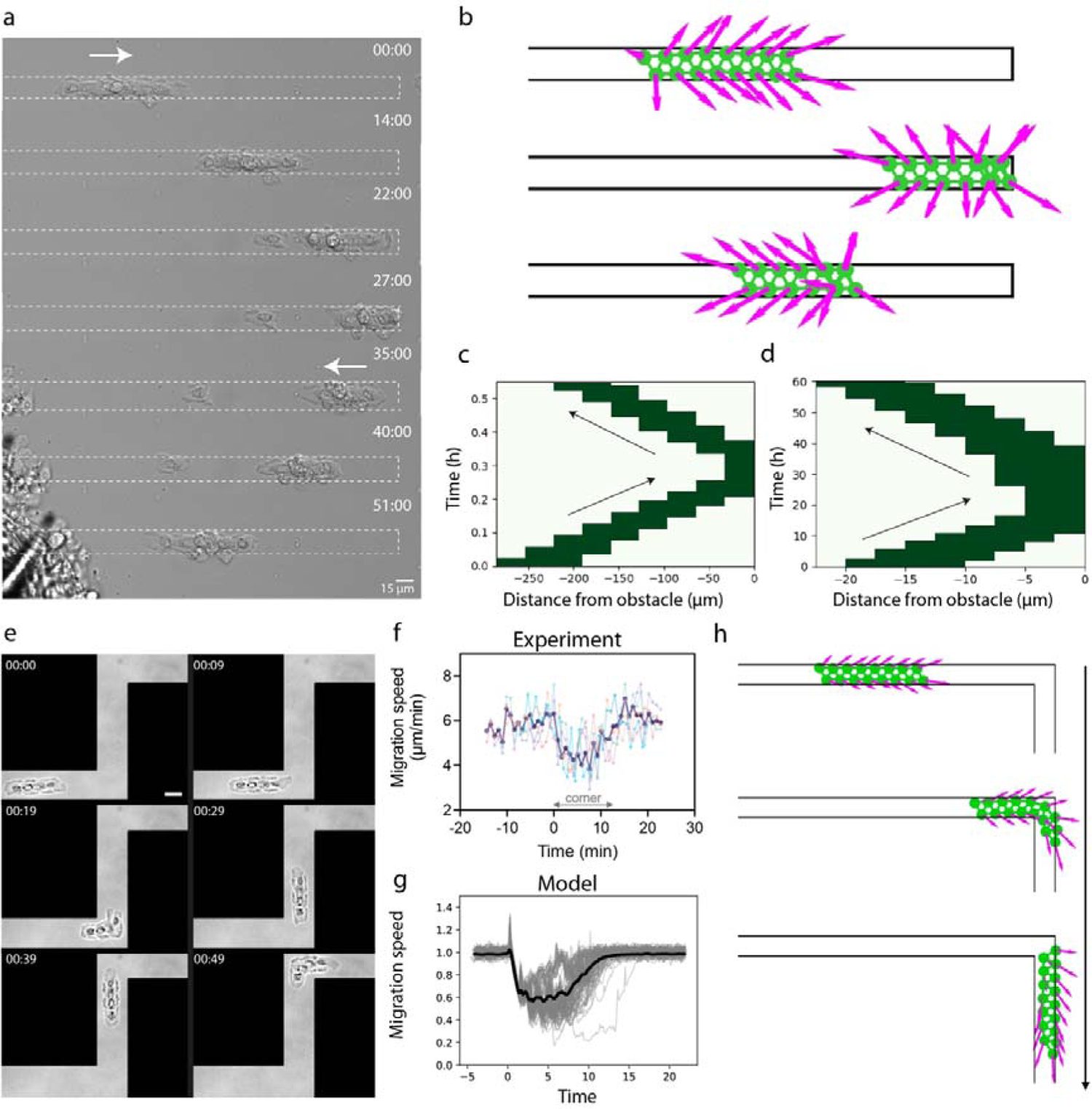
Navigation of cell trains in dead ends and complex microenvironments. **(a)** Time-lapse sequence of the migration of a one-dimensional cluster moving towards the border of a 15 µm-wide fibronectin microstripe (from left to right) during 51 min. After reaching the micropattern extremity on the left part, the cell train compacted against the border, then epithelial cells repolarized, and the cluster migrated in the opposite direction (from right to left). (**b)** Simulation of the migration of a one-dimensional epithelial cluster migrating from the left to the right towards an obstacle. The combination of VA+PA+CRL allows the cell train to repolarize towards the opposite direction after the collision. **(c)** Kymograph of the spatial position over time of the cell train presented in (a), showing its repolarization after the collision. The slopes before and after the collision indicated similar migrating velocities. **(d)** Kymograph of the spatial position over time of the simulated cell train in (b), showing its fast repolarization after the collision and absence of stalling at the boundary. The slopes before and after the collision indicated similar migrating velocities. **(e)** Time-lapse sequence in DIC mode of a cell train migrating on a microstripe of 15 µm wide with corners of 90°. The total duration time is 49 min. Scale bar is 15 µm. **(f)** Superimposed representation of the migration speed of cell trains (L= 4 cells, n=5 from 3 replicates) around corners of 90°. The bold curve in purple represents the mean velocity. **(g)** Predicted speed evolution around a corner of 90°, showing a similar decrease to the experiment, and subsequent recovery of the initial speed. Thin grey curves correspond to simulations of 100 individual clusters, black curve represents the average behavior. **(h)** Time-series of simulated cell clusters navigating around corners of 90°, showing efficient repolarization.

To test the generality of these findings for the collective navigation of cells in complex environments, we further challenged cell cluster migration with geometries including corners of 120° (**Extended Data Fig. 10g**) and 90° angles (**Fig. 5e and Supplementary Movie S16**). Interestingly, we again found experimentally that cell clusters are highly efficient at collectively navigating these complex boundary conditions, with no detected fragmentation despite the induced gradient in velocity orientation. Importantly, the evolution of the migration speed around corners of sharpest angles showed that cell trains only partially and transiently decrease their speed while passing the obstacle (**Extended Data Fig. 10h**). Averaging several experiments on cell trains of L=5 cells showed that they decreased their speed by ∼30% (**Fig. 5f**). These observations are captured by our model, which predicts cell clusters to both remain cohesive and rapidly reorient. Specifically, the model predicts a similar speed decrease to the experimental observation, as well as the fact that cell trains reach their initial speed after a short period of time (**Fig. 5g**), demonstrating their efficient repolarization (**Fig. 5h, Supplementary Movie S17**). Together, these findings not only further support the validity of the combination of VA+PA+CRL, but also point to a potential functional relevance of these specific combination of interactions to allow cell clusters to navigate complex microenvironments.

## Discussion

In summary, in this work we have combined a controlled *in vitro* micropattern approach, force measurement assays and theoretical modelling to dissect the role of contact geometry in migration behavior of small epithelial clusters. We found that not only the size of a cluster, but also its geometry and aspect ratio, was a crucial parameter in determining its migration efficiency, polarization, and state of mechanical stress. We found that axial contacts, established in the direction of cell migration, did not impede migration speed. In contrast, lateral contacts, perpendicular to the direction of cell migration, led to a build-up of lateral polarization, as evidenced by a reorientation of traction forces and monolayer stresses, and thus decreased the migration efficiency of larger cell clusters in proportion to their width. Interestingly, this displays a similar phenomenology, but at the very different length scale of multicellular systems, to how lateral confinement regulates the migration speed of individual cells^40 62^. Our findings are in line with the established importance of intercellular junctions to direct collective cell migration^27 63^, but reveal for the first time their different functional consequences on small cluster migration depending on their geometry relative to the direction of motion. A similar decrease of speed with confinement width has been observed in MDCK monolayers migrating into microchannels, which was explained with the increasing swirling flows and neighbour exchanges and feedback between tissue motility and curvature within the cluster ^19 64^. In contrast, the autonomous clusters we study here do not exhibit such neighbour exchanges on the timescale of cluster migration, placing them in a different class of geometry-dependent migration phenomena.

From a theoretical perspective, we show that while different types of cell-cell interactions give rise to similar large-scale behavior ^36^, introducing anisotropic boundary conditions in small clusters allows us to distinguish them and infer the functional role of each interaction in cluster migration. Indeed, in the presence of boundary conditions, different interaction mechanisms cooperate in complex ways: contact regulation of locomotion (CRL) induces lateral polarization to slow down clusters, polarity alignment (PA) propagates the boundary information from CRL into the bulk; and velocity alignment (VA) breaks symmetry by inducing net polarization in the free axial direction. Thus, these interactions couple the bulk state of stress and polarization in cell clusters to specific boundary conditions. Other cell types such as MDCK have been shown to display flocking behavior experimentally when placed on a ring (periodic boundary conditions)^27^, while we found that cell trains of MDCK cells did not attain a coherent mode of of polarization on open boundary conditions after 6 hours of time-lapse (**Supplementary** Fig. 4), consistent with previous findings on large-scale monolayer expansion^8 59^. This behaviour was predicted by reducing the amplitude of the velocity alignment, leading to a frustrated state with outward polarization for cell trains on microstripes, and a collectively migrating state with coherent polarization in a periodic system (**Supplementary Movie S18**). Interestingly, we found that our model (VA+PA+CRL) captures kerarocyte and MDCK behaviors (**Supplementary Movie S19**), showing more generically how the nature of boundary conditions shape cell cluster behaviors. In the future, it will be interesting to understand at the molecular level the basis for these interactions, whether they are transduced by similar or complementary pathways at the cell-cell contacts, as well as their conservation in other cellular types. This approach could further help advance our understanding of small cluster migration *in vivo*, for instance in the posterior lateral line primordium of zebrafish^6 13^ or in patients with epithelial-originating cancers or carcinomas ^2 14 65^.

## Materials and Methods

### PDMS-coated glass coverslips

Polydimethylsiloxane (PDMS) coated glass coverslips (170 µm thick) were fabricated by spin-coating^66^, as previously described^67^. Briefly, PDMS was prepared from the commercially available Sylgard 184 silicone elastomer kit (Dow Corning, Midland, MI) by mixing the curing agent and the base (1:10 ratio) thoroughly for 2 min. The mixture was degassed, and spin-coated at 5000 rpm on clean 25 mm glass coverslips to obtain a thin PDMS layer of ∼30 µm thick. The PDMS layer was then cured for 3 hours at 60°C, flushed with ethanol and exposed to UV illumination for 15 minutes. PDMS-coated glass coverslips were then stored in the dark at room temperature in a petri dish until use.

### Microcontact Printing

PDMS stamps were used to create micropattern of fibronectin on PDMS-coated glass coverslips. A PDMS mixture (Sylgard 184, Dow Corning Midland, MI) was obtained by mixing the curing agent and the base (1:10 ratio) thoroughly for 2 min^68^. After degassing, the mixture was poured on microstructured silicon wafer which was previously functionalized with fluorosilane (tridecafluoro-1,1,2,2-tetrahydrooctyl-1-trichlorosilane, Gelest) vapors under vacuum to facilitate the removal of the PDMS layer (∼1 cm thick)^69^. After curing overnight at 65°C, the PDMS block was peeled off and cut in stamps of approximately 1 cm^2^. PDMS stamps were washed in an ultrasonic bath with detergent solution (Decon 90, 5%) for 30 min at 35°C, then with isopropanol (70%) for 15 min at 20°C and the stamps were dried under a nitrogen flow A solution was then prepared by mixing 40 µL human fibronectin (Merck) and 960 µL of demineralized water. A volume of 100 µL of this solution was deposited on top of each stamp for 1 h at room temperature in the dark. After gently removing the solution, a PDMS stamp were dried under nitrogen flow and placed carefully in the centre of a PDMS-coated glass coverslip for 15 seconds. PDMS stamps were then gently removed with tweezers and a Pluronic solution at 5 mg/mL was incubated for 5 min at room temperature to passivate non-printed areas. Microprinted PDMS-coated glass coverslips were rinsed three times with sterile PBS and dried under nitrogen flow.

### Cell culture

Fish epithelial keratocytes were obtained from the scales of Central American cichlid (*Hypsophrys Nicaraguensis*)^40 67^. Scales were gently taken off the fish and placed in the center of a microprinted PDMS-coated glass coverslips and covered with a drop of 150 µL of culture medium. The culture medium was composed of Leibovitz’s L-15 medium (Thermo Fisher Scientific) supplemented with 10% fetal bovine serum (FBS, Capricorn), 1% penicillin/streptomycin (Westburg), 14.2 mM HEPES (Sigma Aldrich) and 30% deionized water were put on top of the scale. A glass coverslip of 22 mm in diameter was deposited on top of the scales and few drops of culture medium were added around the samples. Epithelial keratocytes were cultured in the dark at room temperature for 12 h.

Epithelial cells from the Madin-Darby Canine Kidney cell line (MDCK II, Sigma #85011435) were maintained in polystyrene T75 flasks of in a cell culture incubator at 37°C and 5% CO_2_. MDCK cells were cultured in proliferation medium composed of Dubelcco’s Modified Eagle’s medium (DMEM), high glucose (4.5 g/l) with L-glutamine (BE12-604F, Lonza) supplemented with 10% (v/v) Fetal Bovine Serum (FBS, AE Scientific) and 1% of penicillin and streptomycin antibiotics (AE Scientific).

### Time-lapse imaging

Time-lapse microscopy experiments were carried out on a Nikon Ti-U inverted microscope (Nikon, Japan) equipped with a manual stage^40 67^. Differential Interference Contrast (DIC) images were taken every 3 min using a ×10, ×20 or ×40 objective and captured with a DS-Qi2 camera (Nikon, Japan) controlled with the NIS Elements Advanced Research 4.0 software (Nikon). Tracking of the cell clusters were performed with the Manual Tracking plugin on FIJI.

### Drug treatment

EGTA (Sigma Aldrich) was added to the normal medium at a final concentration of 2 mM to partially chelate the calcium in the medium.

### Mitochondrial membrane potential

The mitochondrial membrane potential was measured using MitoTracker (MT) Red that stained active mitochondrial in live cells by binding thiol-reactive chloromethyl groups in the mitochondrial membrane. A concentration of 50 nM of the MitoTracker Red dye (Invitrogen) was used for 30 min at room temperature to allow dye equilibration across the plasma and inner mitochondrial membranes. For imaging, the medium containing the MitoTracker Red dye was replaced with fresh normal medium.

### Immunofluorescence and confocal microscopy

Fish keratocytes were fixed and permeabilized with 4% paraformaldehyde and 0.2% Triton X-100 in PBS for 15 min at room temperature. The samples were then incubated with 1% BSA in PBS for 30 min at room temperature, followed by sequential incubation with primary and secondary antibodies diluted with 1% BSA in PBS for 45 min at 37°C. Actin filaments were stained with AlexaFluor 488 phalloidin (Invitrogen, 1:200), the nucleus with 4′,6-diamidino-2-phenylindole (DAPI; Invitrogen, 1:200), microtubules with an anti-tubulin antibody produced in mouse (1:200) and the Golgi apparatus with 10 μg/ml wheat germ agglutinin (WGA) conjugated with Alexa Fluor 594. Images were collected in epifluorescence and confocal mode with a Nikon A1R HD25 motorized inverted microscope equipped with x20, x40, x60 Plan Apo (NA 1.45, oil immersion) and x100 Plan Apo silicone objectives and lasers that span the violet (405 and 440 nm), blue (457, 477, and 488 nm), green (514 and 543 nm), yellow-orange (568 and 594 nm), and red (633 and 647 nm) spectral regions. Epifluorescence images were recorded with a photometrics Prime 95B camera (Photometrics Tucson, AZ) using NIS Elements Advances Research 4.5 software (Nikon). Confocal images were recorded with x100 Plan Apo silicone objective of high numerical aperture (Plan Apochromat Lambda S 100x Silicone, Nikon Inc) in galvanometric mode with small Z-depth increments (0.1 μm) and a pinhole of 12 µm to capture high resolution images. Confocal images were processed using NIS-Elements (Nikon, Advanced Research version 4.5).

### Polyacrylamide hydrogels

Polyacrylamide gel substrates of 18 kPa were prepared as previously described^70 71^. Briefly, glass-bottom dishes (MatTek, 35 mm) were treated with a solution of 714 µL acetic acid (%), 714 µL of silane in 10 mL of ethanol (96%) for 20 min. After the removal of the solution, the dishes were rinsed twice with ethanol (96%) and dried under nitrogen flow. For 18 kPa gels, a solution of 0.65 mg of N-hydroxyethyl acrylamide (HEA) in 5 mL of PBS was prepared. 820 µL of this solution was mixed in a microcentrifuge tube (Eppendorf, 1.5 mL) with 100 µL bisacrylamide (2%), 80 µL acrylamide (40 %) and 3 µL red fluorescent carboxylate-modified beads (0.2 µm, red 580/605, Life Technologies). Degas under vacuum for at least 20 min and then 0.5 µL of N,N,N’,N’-tetramethyl ethylenediamine (TEMED) with 5 µL of ammonium persulfate (APS) were added to initiate the polymerization. 25 µL of the solution was carefully put on each glass-bottom dishes and 18 mm glass coverslips (previously treated under corona) were placed on top of them. After 1 h polymerization under oxygen-free atmosphere, the coverslips were removed from the gels in hot water by using tweezers. PDMS stamps were then used to create micropattern of fibronectin on each gel. After washing the stamps in an ethanol solution (70%) for 15 min in an ultrasonic bath and drying under nitrogen flow, a Corona was used for 30 s to make them more hydrophilic. A solution was prepared by mixing 70 µL human plasma fibronectin (Merck), 20 µL green fibrinogen and 910 µL PBS. 100 µL of this solution was put on top of each stamp for 1 h at room temperature in the dark. After the removal of the solution, the stamps were dried under nitrogen flow and deposited carefully in the center of each gel. We pressed carefully on each corner of the stamp with tweezers to allow protein transfer. The printing process occurred overnight at 4°C in the dark. Stamps were then gently removed with PBS and gels were rinsed 3 times with PBS. Bovine serum albumin (BSA) solution (5 mg/mL) was then used for 4 hours at and 4°C to passivate areas without any protein on the gels followed by 3 rinses with PBS. Cells were then seeded on microprinted hydrogels.

### Traction force microscopy

Fluorescent beads of 200 nm in diameter were homogeneously added in the polyacrylamide solution before polymerization and their position was imaged over time with an IX83 inverted microscope equipped with a 632/22 excitation filter (incorporated in the Spectra-X light engine), a glass dichromatic mirror (Olympus), and FF01-692/40-25 (Semrock) emission filter. A reference image was obtained after the removal of cells by trypsinization. The 2D displacement field was computed using a custom PIV implementation in Matlab (MathWorks). We used Fourier-transform traction microscopy (FFTM)^59 72 73^ and filtered the displacement field accordingly implementing a predictor-corrector filter approach. Criterions for filtering were selected through semi-quantitative approaches to ensure that noise in cell-free areas is acceptable when compared to the cell tractions. Experimental and digital noise in the displacement field were minimized using a high density of fiducial markers, an appropriate size and overlap of the PIV interrogation window, and an algorithm to avoid peak-locking effects ^59^. Noise in the measured displacement field was reduced by implementing a predictor-corrector filter approach, as described in ^74^. We used a combination of filter power z=0.01 and kernel size of 2.

### Monolayer stress microscopy

We employed the monolayer stress microscopy (MSM) method^61^ to quantify the spatial distribution of intracellular stress in cell clusters. This method is based on force equilibrium between cell-substrate tractions and cell-cell stresses. Implementation of MSM starts with the recovery of local tractions exerted by the monolayer upon its substrate. We consider a monolayer of contiguous cells which forms a uniform sheet that is flat and thin, meaning that, compared with the lateral span of the monolayer, its height is negligible. In that case, stresses within the monolayer and underlying tractions exerted by the monolayer upon its substrate are taken to be planar with no out-of-plane contributions. The stresses everywhere in the monolayers are then determined by terms arising from tractions and boundary conditions. Since the monolayer always remains in mechanical equilibrium, Newton’s laws demand that this force balance must be independent of material properties of the monolayer. MSM was implemented, as a custom-made software, in Python 3 using NumPy^75^, SciPy^76^, Matplotlib^77^, scikit-image^78^, pandas^79^, pyFFTW^80^, opencv^81^ and cython^82^.

### Statistical analysis

Each experiment was repeated at least three times. Every set of data was tested for normality test using the d’Agostino-Pearson test in Prism 10.0 (GraphPad) that combines skewness and kurtosis tests to test whether the shape of the data distribution was similar to the shape of a normal distribution. For paired comparisons, significances were calculated in Prism 10.0 (GraphPad) with a Student’s *t*-test (two-tailed, unequal variances) when the distributions proved to be normal. If a data set did not pass the normality tests, the significances were calculated with Mann–Whitney (two-tailed, unequal variances). For multiple comparisons with non-normal distribution, data sets were analyzed with a Kruskal-Wallis test in Prism 10.0 (GraphPad), which is a suitable nonparametric test for comparing multiple independent groups when the data are skewed. When the null hypothesis was not retained (p-value < 0.05), Kruskal-Wallis was corrected with Dunn’s test, which is a nonparametric test with no pairing and multiple comparisons that can be used for both equal and unequal sample sizes. Unless otherwise stated, all data are presented as mean ± standard deviation (s.d.). The confidence interval in all experiments was 95% and as a detailed description of statistical parameters it is included in all figure captions with *p < 0.05, **p < 0.01, ***p < 0.001, ****p<0.0001 and n.s. is not significant.

### Ethical compliance

The use of primary epithelial cells, keratocytes, harvested from the scale of *hypsophrys nicaraguensis* was done in accordance with European guidelines for animal experimentation and with the agreement of the local ethic committee of the University of Mons which has reviewed the procedure.

### Data availability

The data that support the plots within this paper and other finding of this study are available from the corresponding authors upon request.

## Supporting information

Supplementary Theory Note

## Acknowledgments

M.L., E.V. and S.G. acknowledges funding from FEDER Prostem Research Project no. 1510614 (Wallonia DG06), the F.R.S.-FNRS Epiforce Project no. T.0092.21, the F.R.S.-FNRS Cellsqueezer Project no. J.0061.23, the F.R.S.-FNRS Optopattern Project no. U.NO26.22 and the Interreg MAT(T)ISSE project, which is financially supported by Interreg France-Wallonie-Vlaanderen (Fonds Européen de Développement Régional, FEDER-ERDF). A.R. is financially supported by F.R.S.-FNRS as Research Fellow (ASP). E.V. and Y.K. are financially supported by F.R.S.-FNRS as FRIA Grantees. This project was supported by the European Research Council under the European Union’s Horizon 2020 Research and Innovation Program Grant Agreements 851288 (to E.H) and the Marie Skłodowska-Curie grant agreement 797621 (to M.G.G.). D.B.B. was supported by the NOMIS foundation as a NOMIS Fellow and by an EMBO Postdoctoral Fellowship (ALTF 343-2022), and performed this work in part at Aspen Center for Physics, which is supported by National Science Foundation grant PHY-1607611. X.T. and M.G.G. acknowledge support from The Generalitat de Catalunya (AGAUR SGR-2017-01602 to and the CERCA Programme), the Spanish Ministry for Science and Innovation MICCINN/FEDER (PGC2018-099645-B-I00), the European Research Council (Adv-883739), Fundació la Marató de TV3 (201903-30-31-32), the European Commission (H2020-FETPROACT-01-2016-731957), La Caixa Foundation, and by CIBER -Consorcio Centro de Investigación Biomédica en Red-(CB15/00153), Instituto de Salud Carlos III, Ministerio de Ciencia e Innovación. IBEC is recipient of a Severo Ochoa Award of Excellence from the MINECO.

## Author contributions

S.G. and E.V. conceived the project and S.G. supervised the project. E.V. developed minimal epithelial models and performed cell experiments, tracking and imaging with A.R. D.B.B. and E.H. developed the theoretical model, D.B.B. implemented and performed simulations. A.R., L.R., Y.K. and M.L. contributed to experiments. E.V., D.B.B., A.R., M.G.G., X.T., E.H. and S.G. analyzed data. The article was written by E.V., D.B.B., E.H. and S.G., read and corrected by all authors, who all contributed to the interpretation of the results.

---

**Extended Data Figure 1.**
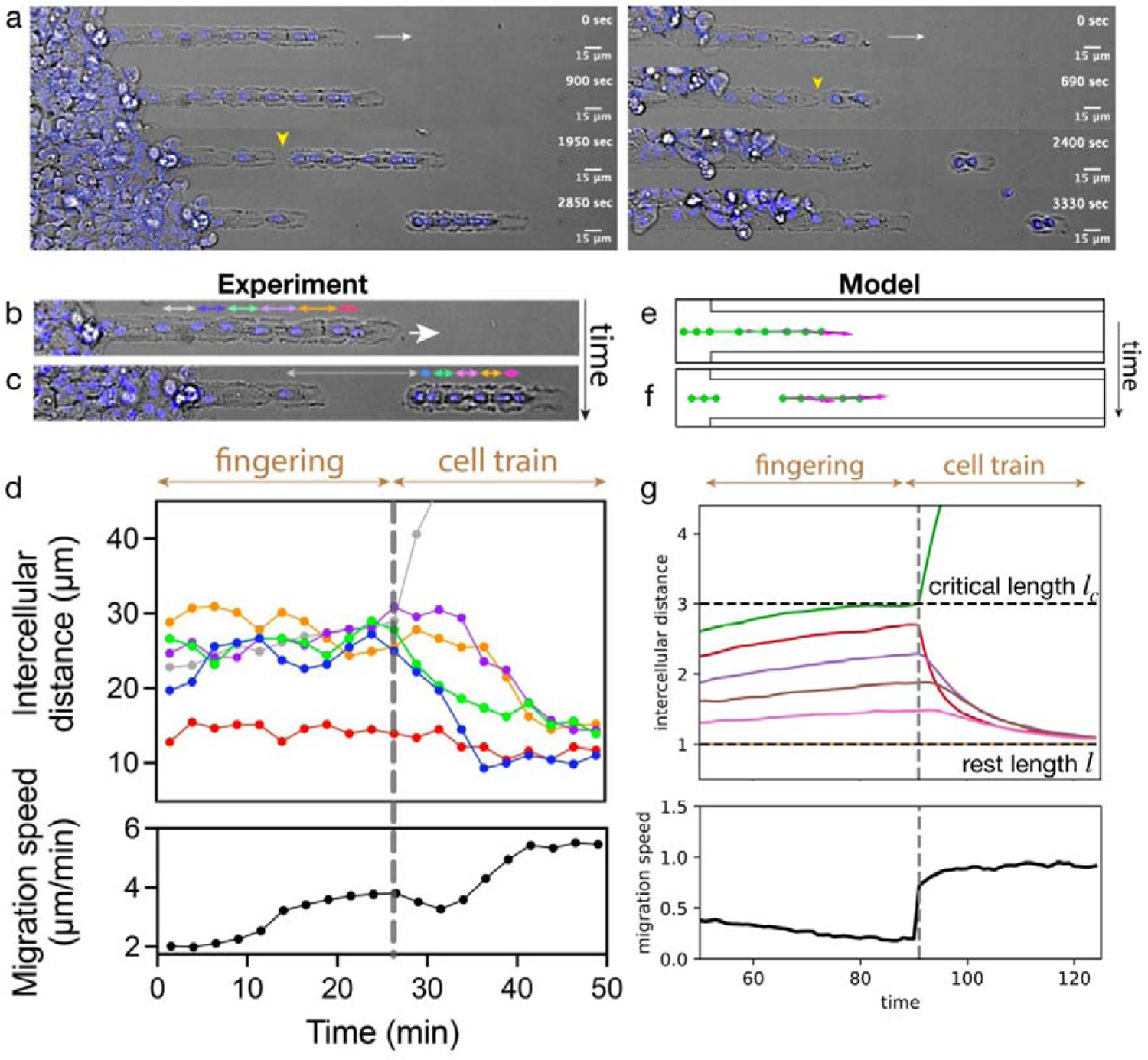
Digitation of a primary epithelial tissue and fragmentation into cell trains. **(a)** Time-lapse sequences in Differential Interference Contrast (DIC) mode of the formation of cell trains of L=6 cells (left) and L=2 cells (right) from the fragmentation of monolayer extensions formed on 15 µm wide micropatterns. Nuclei are stained in blue with Hoechst 33342. White arrow shows the direction of migration and yellow arrows the fragmentation sites. Scale bars are 15 µm. **(b)** Time lapse sequence in DIC mode of the extension of a “finger” of epithelial keratocytes on a 15 µm wide microstripe, which remained attached to the cell mass. Nuclei are stained in blue with Hoechst 33342. White arrow shows the direction of migration, while arrows in red, orange, purple, green and blue and yellow represent the intercellular distance (*l*) between five first individual cells within the tissue extension. The grey arrow corresponds to the intercellular distance at the fragmentation site. Intercellular distances are determined by the distance between nuclei. **(c)** At severe stretching, the tissue extension fragments into an autonomous cell train. Scale bars are 15 µm. **(d)** Evolution of intercellular distances versus time during the fingering process and after the fragmentation (grey dashed line). The intercellular distances are color-coded to their position from front to rear: red, orange, purple, green and blue. The grey dots represent the intercellular distance at the fragmentation site. The migration speed of the leading front is indicated in black. **(e-g)** Equivalent plots for simulations of the fingering process, which back cells simulated to be slowly moving. The initial fragmentation process allows to infer a critical length of the cell-cell contacts, which is estimated as *l*_c_ ≍ 3*l* (see SI Theory Note for details).

**Extended Data Figure 2.**
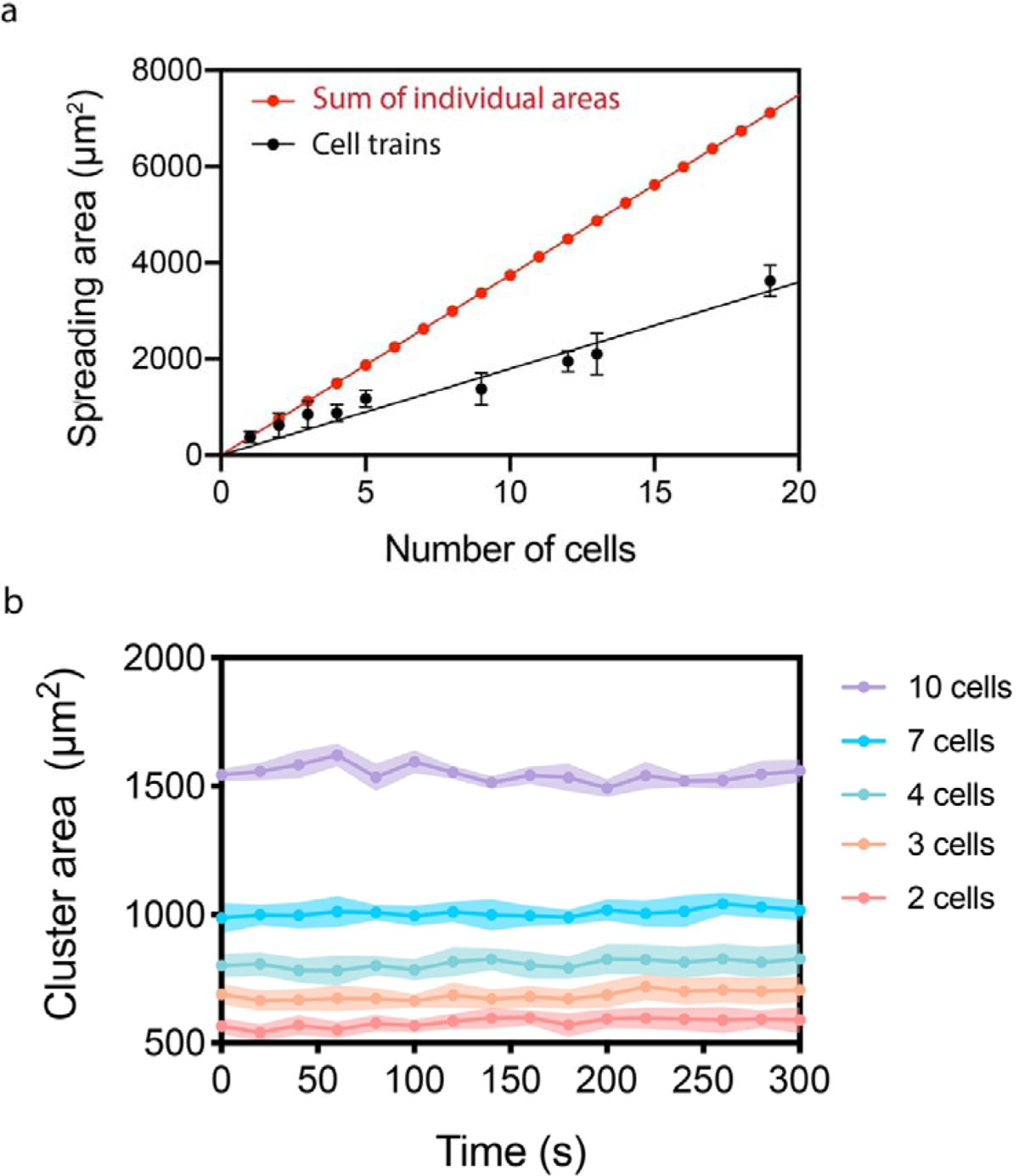
Autonomous cell trains are compacted and cohesive units. **(a)** The cluster area is linearly correlated with the number of cells (black points) and smaller than the sum of individual cell areas (red points), suggesting that cells in one-dimensional epithelial clusters are significantly compacted. **(b)** The cluster area is constant over time, regardless the number of cells, demonstrating that one-dimensional epithelial clusters move as a single and cohesive unit. Error bars denote standard deviations.

**Extended Data Figure 3.**
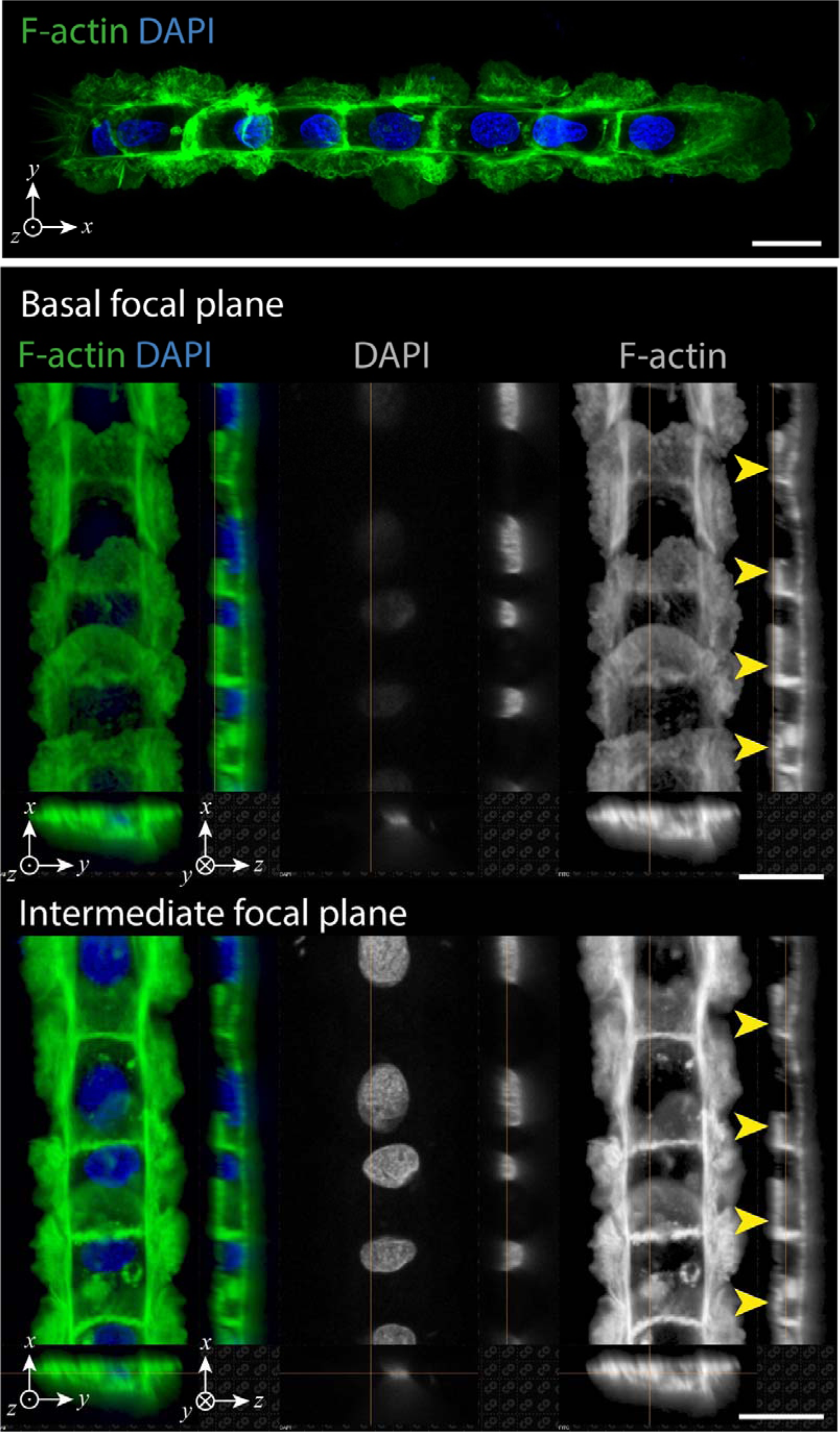
Cell trains are composed of cryptic lamellipodia that invade underneath the adjoining cell. **(a)** Normal view in high resolution confocal mode of a cryptic lamellipodia in a cell train of 15 µm wide composed of 8 cells. **(b)** Zoomed side views of basal and intermediate focal planes indicating the presence of cryptic lamellipodia (yellow arrows) that extend underneath the cell body of the preceding cell. Actin is labelled in green with Phalloidin and DNA in blue with DAPI. The white arrows show the position of typical cryptical lamellipodia. Scale bars are 15 µm.

**Extended Data Figure 4.**
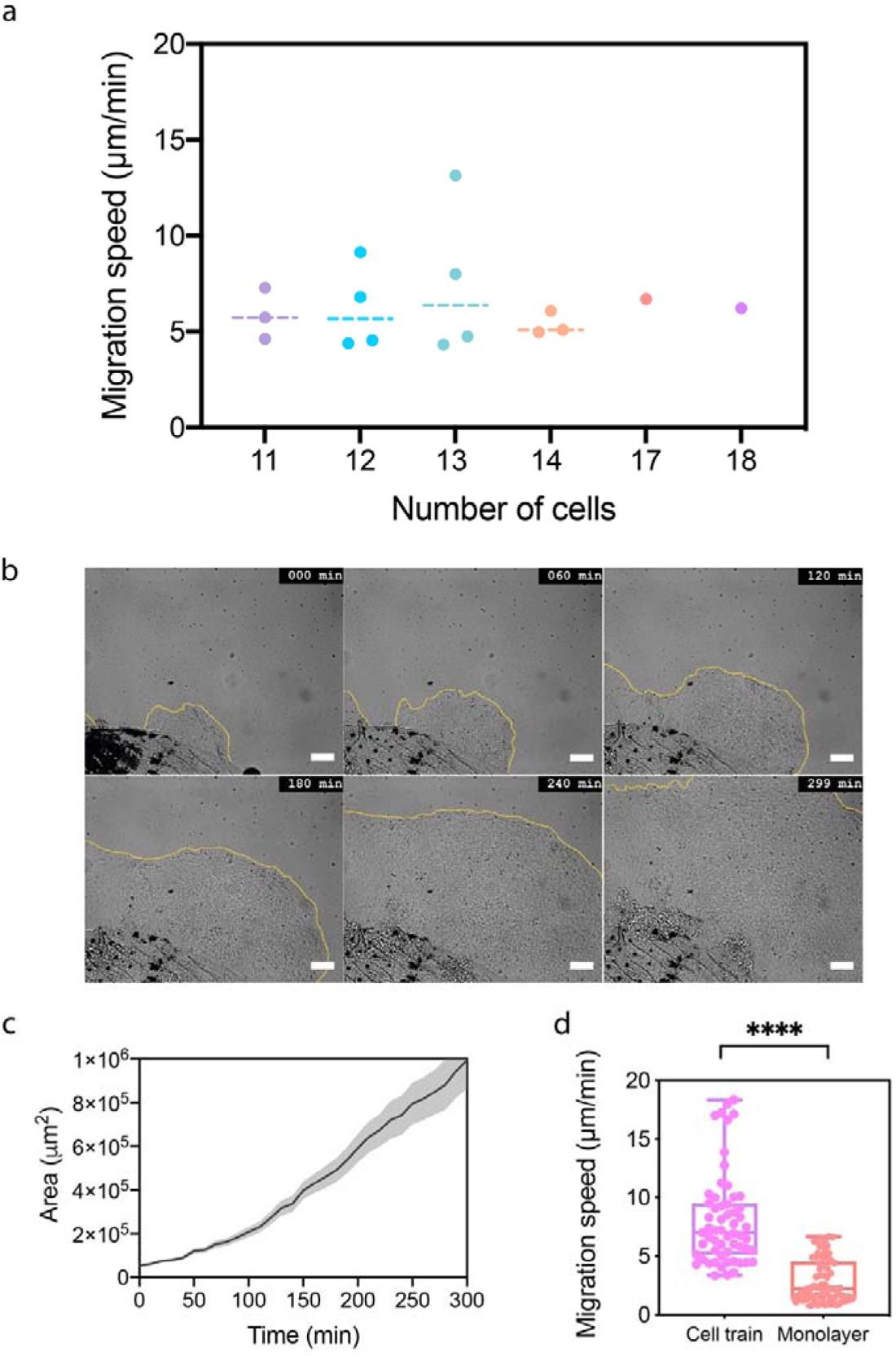
Migration of very long autonomous cell trains and extension of a primary epithelial monolayers. **(a)** Migration speed of very long cell trains. Each point represents the mean migration speed of one cell train, whose length ranges from 11 to 18 cells. Dashed lines represent the median. The migration speed was calculated from time-lapse experiments of 45 min. The total number of cell trains is 16. **(b)** Representative time-lapse sequence in DIC mode of an epithelial monolayer of primary keratocytes growing out of a fish scale during 299 min. The growing front is depicted with a yellow line. The scale bar is 100 µm. (c) Evolution of the monolayer area over time (n=3, mean ± S.D.). (d) Migration speed for one-dimensional clusters (“cell train” in pink, n=62) and epithelial monolayers (“monolayer” in light red, n=74) with ****p < 0.0001.

**Extended Data Figure 5.**
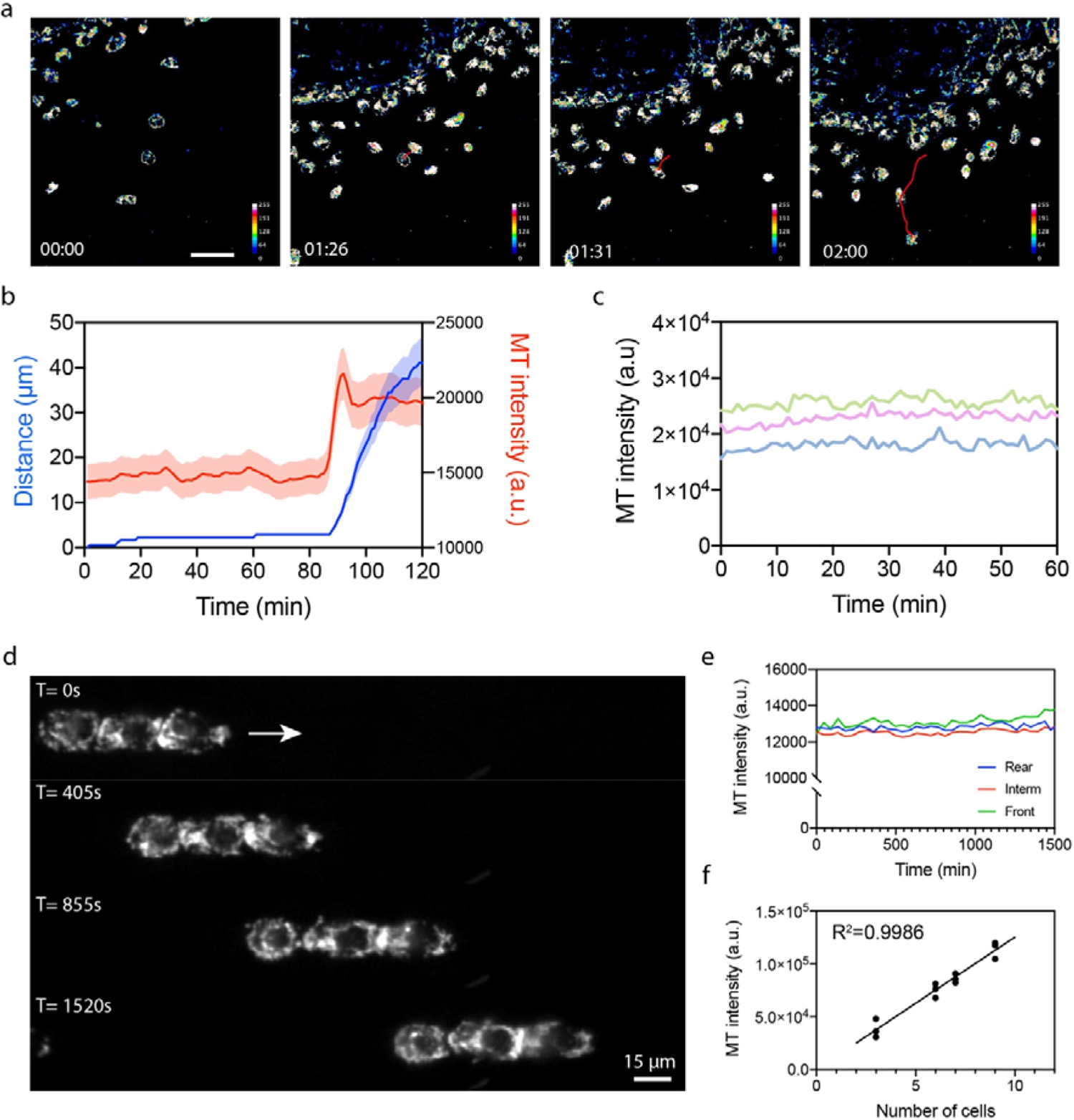
The mitochondrial membrane potential increases as the cell polarizes and remains constant during migration. **(a)** Typical time-lapse sequence in epifluorescence mode of single epithelial cells stained with red MitoTracker (MT) that stained active mitochondrial in live cells by binding thiol-reactive chloromethyl groups in the mitochondrial membrane. Cells were digitized in 256 bits and the MitoTracker intensity was color-coded (from high to low: white, purple, red, orange, yellow, green, light blue and dark blue). The redline shows the displacement of a cell that started to polarize and migrate at 86 min. The scale bar is 30 µm. **(b)** Evolution of the distance (in blue) and the MT intensity (in red) over time (n=3, mean ± S.D.) **(c)** Evolution of the MT intensity of three individual polarized cells that migrate during 60 min. **(d)** Typical time-lapse sequence of a cell train (L=3 cells) migrating on a fibronectin microstripe of 15 µm wide and labeled in live conditions with mitotracker red. The total duration is 1520 sec. and the scale bar is 15 µm. (e) Temporal evolution of the mitochondrial potential membrane intensity per cell in cell trains of L=3 cells (n=6 cell trains). **(f)** Linear evolution of the mitotracker intensity for cell trains of L=3, 6, 7 and 9 cells (n=3 for each condition) with R^2^=0.9986.

**Extended Data Figure 6.**
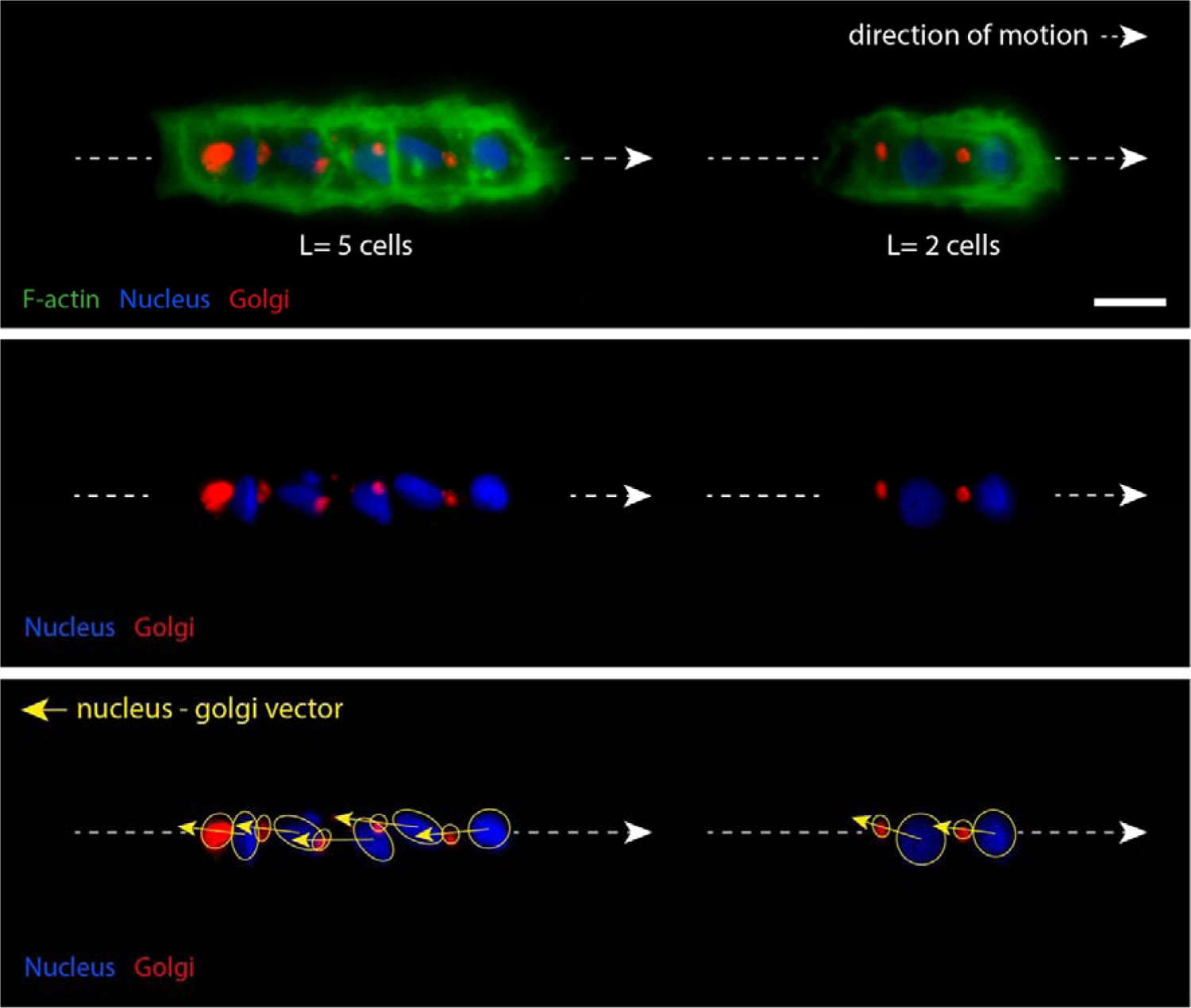
The Golgi complex is positioned behind the nucleus in cell trains. Epifluorescent images of trains of cells composed of L=5 (left) and L=2 (right) cells migrating on fibronectin microstripes of 15 µm wide. Keratocytes were labelled for the F-actin (green), the Golgi complex (red) and the nucleus (blue). The direction of motion is depicted by a white arrow and the orientation of the Golgi complex relative to the nucleus is indicated by a yellow arrow. The white dashed line represents the cell train axis. The scale bar is 15 µm.

**Extended Data Figure 7.**
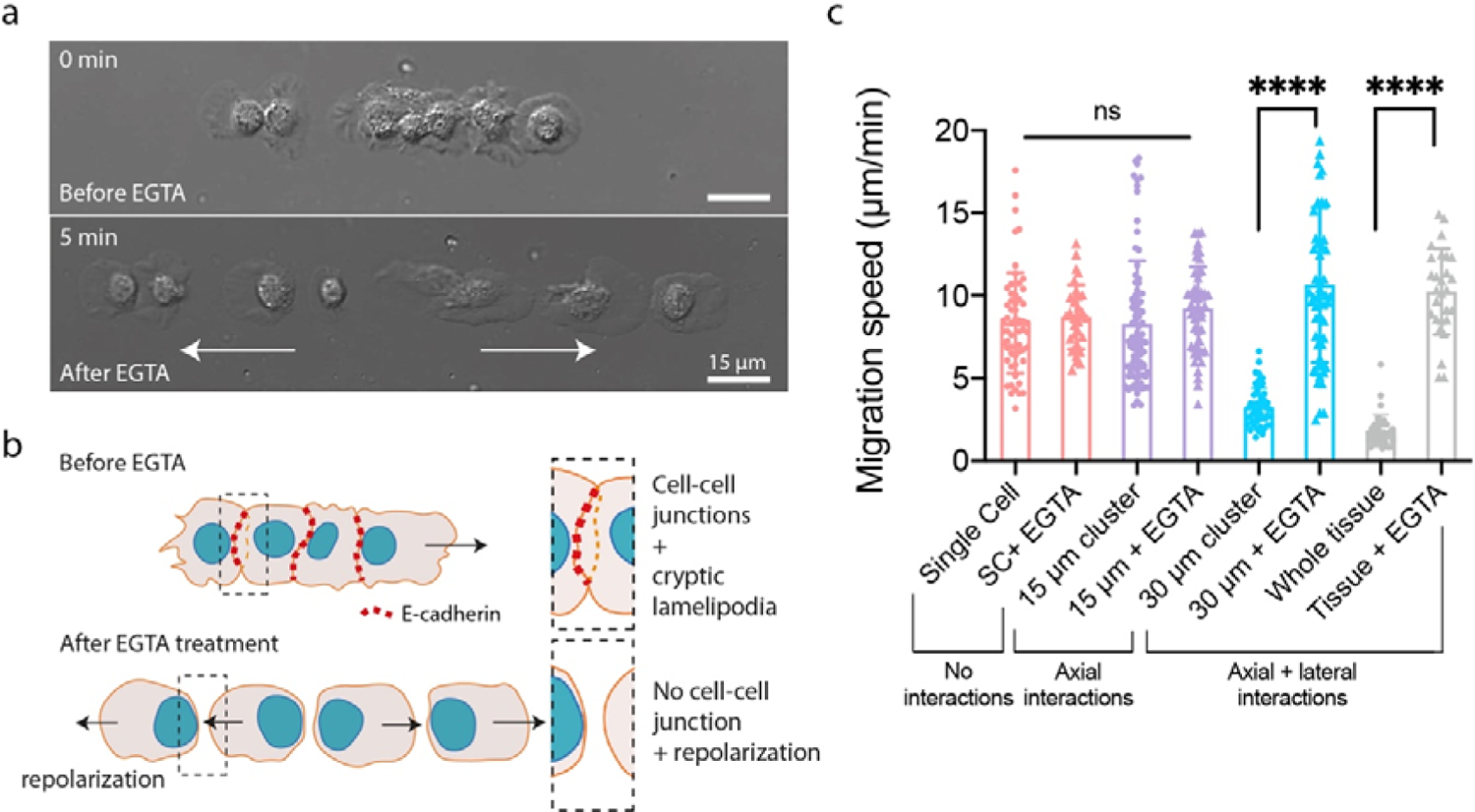
Disruption of cell-cell adhesions with EGTA treatments leads to the increase of the migration speed and repolarization events. (a) Typical microscopy images in DIC mode of a one-dimensional epithelial cluster migrating on a 15 µm wide micropatterns before (t=0 min) and after (t=5 min) EGTA treatments. White arrows show the cell repolarization. Scale bars are 15 µm. (b) Schematic representation of the EGTA effect that disrupts adherens cell-cell adhesions due to a weakening of the rigid extracellular domain of E-cadherin and leads to repolarization events. (c) Migration speed of single cells on 15 µm wide microstripes (control: n=57, EGTA: n=32, in red), cell trains on 15 µm wide microstripes (control: n=88, EGTA: n= 49, in purple), epithelial clusters on 30 µm wide microstripes (control: n=51, EGTA: n=57, in blue) and whole epithelial monolayers (control: n=37, EGTA: n=28, in grey). Control data are circles and EGTA-treated data are lozenges, ns is non-significant and ****p < 0.0001.

**Extended Data Figure 8.**
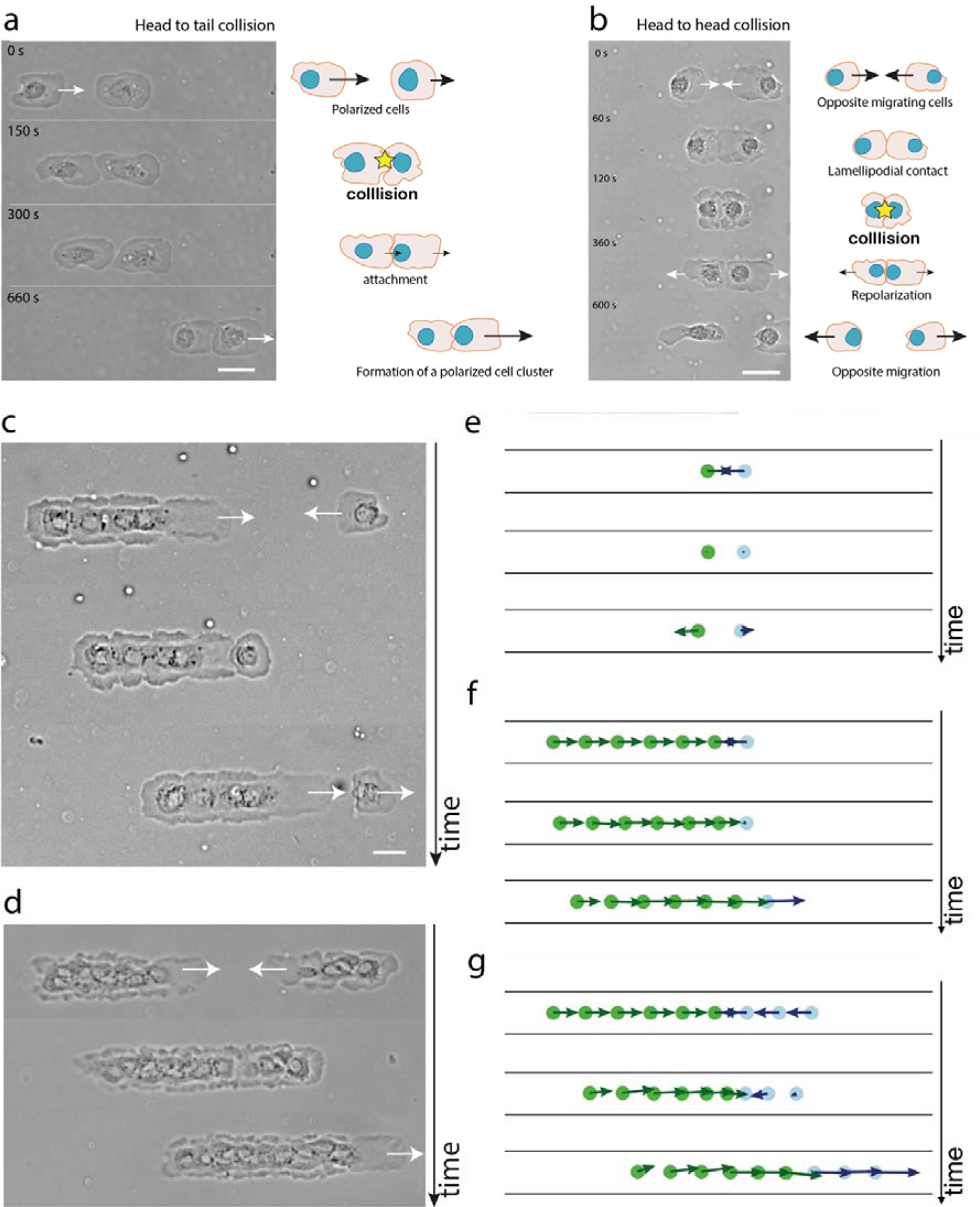
Different collision scenarii between single epithelial cells and cell clusters. The collision configuration between two single epithelial cells induces an attractive or repulsive response. **(a)** Head-to-tail collisions promote the formation of a contact between a cell lamellipodia and the tail of a neighbouring cell, leading to the formation of a polarized cell doublet. (b) Head-to-head collisions between individual epithelial cells show a repulsive response of both cells that repolarized rapidly in opposite directions. Collisions events between **(c)** on single cell against a cell train and **(d)** two cell trains always lead to the repolarization of the smaller cluster. Simulations of collision between **(e)** two opposite single cells, **(f)** on single cell against a cell train and **(g)** two cell trains show a repulsive response of both individual cells that repolarized rapidly in opposite directions and the repolarization of the shorter cluster, as observed experimentally in (b-d).

**Extended Data Figure 9.**
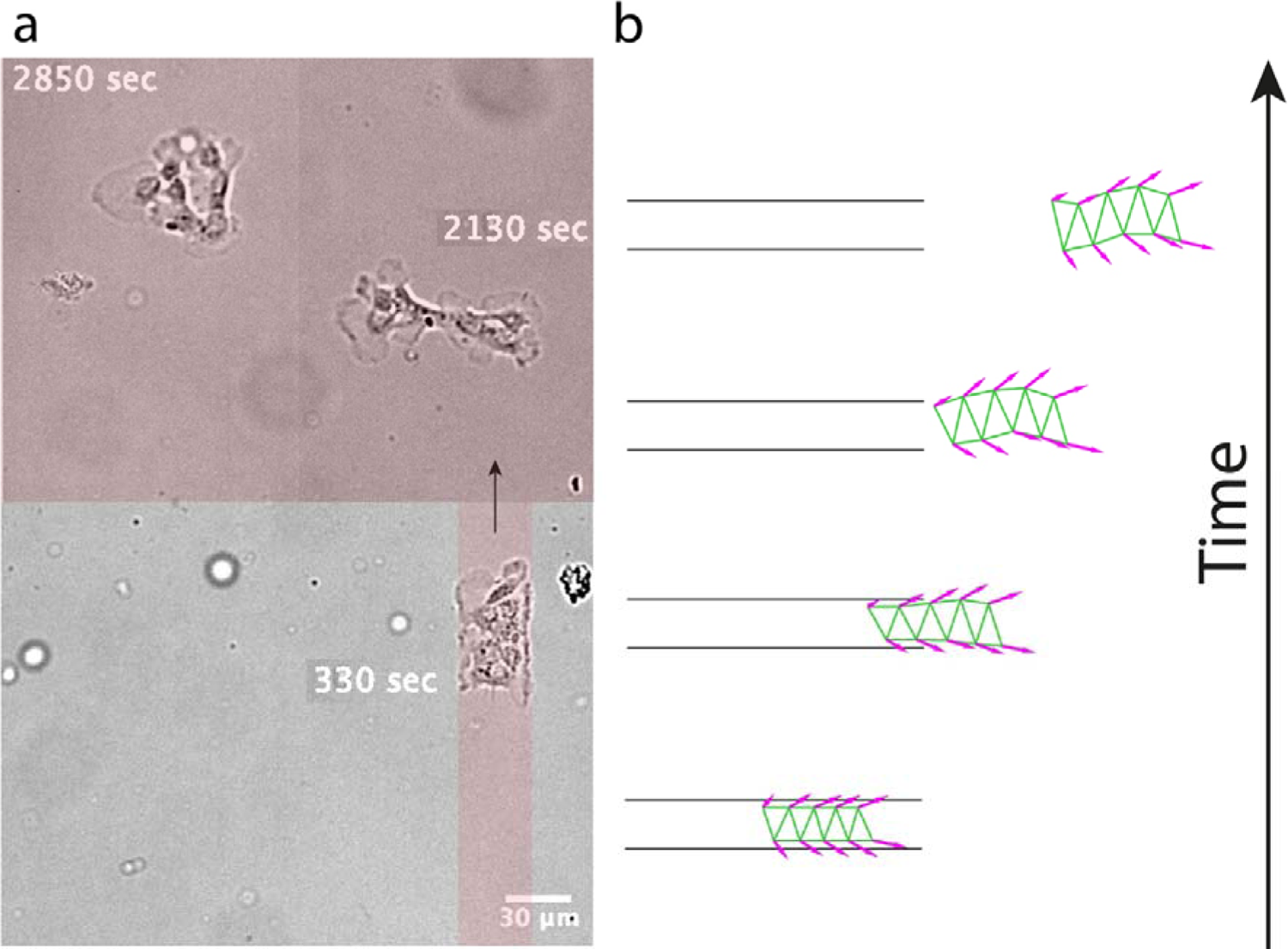
Unconfined migration of a cell cluster. **(a)** Time-lapse sequence in differential Interference Contrast (DIC) mode of a cell cluster reaching the end of a 30 µm wide adhesive microstripe and entering a free open adhesive space. Upon exiting the microstripe, the cell cluster rapidly develops lamellipodia in the lateral directions away from their neighbours and finally breaks apart. The black arrow shows the direction of migration, while the light red zones indicate the microprinted adhesive areas. **(b)** Theoretical simulation of the unconfined migration of a one-dimensional epithelial cluster migrating out of a confined zone from the left to the right.

**Extended Data Figure 10.**
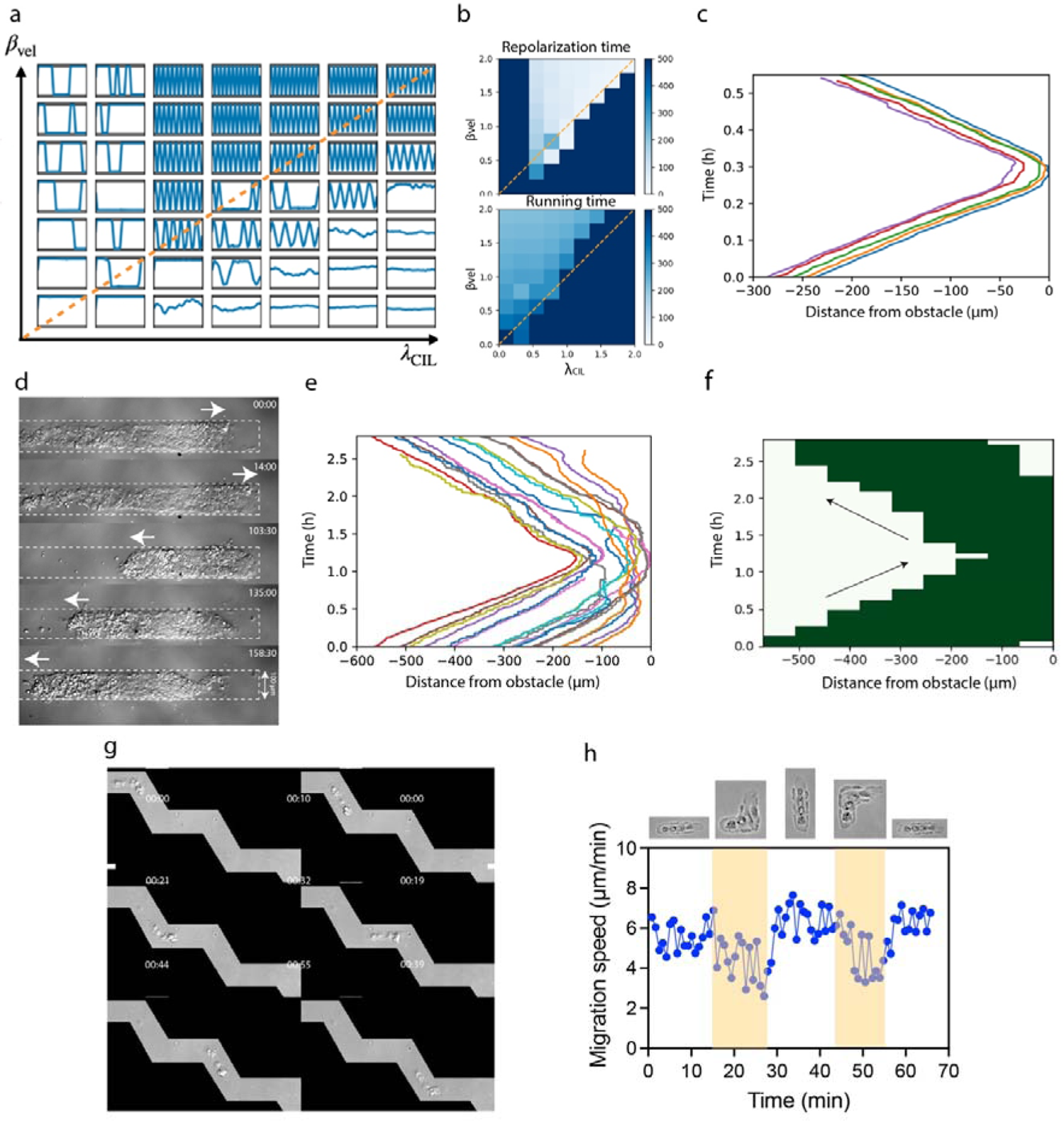
Perturbation of the polarization of epithelial cell clusters by migrating against an obstacle and in complex turns. **(a)** Phase diagram showing trajectories of cell cluster center of mass in obstacles at both ends of the microstrip, as a function of the interaction parameters β_vel_ and β_CRL_. **(b)** Phase diagrams of repolarization and running times as a function of β_vel_ and β_CRL_, quantifying the repolarization and running phases in panel (b). **(c)** Distance travelled over time by individual cells within the cell train moving towards the border of a 15 µm-wide fibronectin microstripe (from left to right) during 51 min. After reaching the micropattern extremity on the left part, the cell train compacted against the border, then epithelial cells repolarized, and the cluster migrated in the opposite direction (from right to left). Each color-coded line corresponds to one cell. **(d)** Time-lapse sequence of the migration of a 100 µm wide epithelial cluster moving towards the border of a fibronectin microstripe (left to the right) during 158 min. After reaching the micropattern extremity on the left part, the wide cluster compacted against the border, then epithelial cells repolarized and the whole cluster migrated in the opposite direction (right to left). **(e)** Distance travelled over time by individual cells within the 100 µm wide cluster presented in (d). Each color-coded line corresponds to the one cell. **(f)** Kymograph of the spatial position over time of the 100 µm wide cluster presented in (d), showing its repolarization after the collision. The slopes before and after the collision indicated similar migrating velocities. **(g)** Time-lapse sequence in DIC mode of a cell train migrating on a microstripe of 15 µm wide with corners of 120°. The total duration time is 55 min and scale bars are 15 µm. **(h)** Representative example of the decrease of the instantaneous velocity of a cell train (L= 4 cells) around corners of 90°.

**Supplementary Movie S1 –** Time-lapse sequence in Differential Interference Contrast (DIC) mode of the formation of cell trains of L=2 cells from the fragmentation of a tissue extension formed on 15 µm wide micropattern. Nuclei are stained in blue with Hoechst 33342. The scale bar is 15 µm.

**Supplementary Movie S2 –** Time-lapse sequence in Differential Interference Contrast (DIC) mode of the formation of cell trains of L=5 cells from the fragmentation of a tissue extension formed on 15 µm wide micropattern. Nuclei are stained in blue with Hoechst 33342. The scale bar is 15 µm.

**Supplementary Movie S3 –** Time-lapse sequence in DIC mode of a polarized one-dimensional cluster (L=7 cells) migrating on a 15 µm wide fibronectin micropattern. The scale bar is 10 µm.

**Supplementary Movie S4 –** Time-lapse sequence in DIC mode of a polarized one-dimensional cluster (L=10 cells) migrating on a 15 µm wide fibronectin micropattern. The scale bar is 10 µm.

**Supplementary Movie S5 –** High resolution confocal Z stack of cryptic lamellipodia in a cell train of 15 µm wide composed of L=8 cells. Cryptic lamellipodia extend underneath the cell body of the preceding cell. F-actin is stained in green with Phalloidin.

**Supplementary Movie S6 –** Time-lapse sequence in epifluorescence mode of epithelial keratocytes labelled with the MitoTracker (MT) Red dye. Cells were detached from the front edge of primary monolayer growing from the fish scale. The fluorescence intensity of the dye was proportional to the mitochondrial membrane potential. Unpolarized and stationary cells showed a low MT fluorescence intensity, which increased immediately with their polarization and migratory state. Cells were digitized in 256 bits and the MT intensity was color-coded (from high to low: white, purple, red, orange, yellow, green, light blue and dark blue). The total duration time is 1 hour and 59 minutes.

**Supplementary Movie S7 –** Time-lapse sequence in epifluorescence mode of a cell train of length L=3 cells labelled with the MitoTracker (MT) Red dye. The total duration is 1520 seconds and the scale bar is 15 µm.

**Supplementary Movie S8 –** Time-lapse sequence in DIC mode of a polarized epithelial cluster that migrates on a 35 µm wide fibronectin micropattern. The scale bar is 25 µm.

**Supplementary Movie S9 –** Time-lapse sequence in DIC mode of a polarized epithelial cluster that migrates on a 50 µm wide fibronectin micropattern. The scale bar is 25 µm.

**Supplementary Movie S10 –** Animated simulations of the model with single interaction types; from top to bottom: polarity alignment (PA), velocity alignment (VA), stress-polarity coupling (SPC), contact regulation of locomotion (CRL). Green dots: cell positions, green lines: elastic links between cells, magenta arrows: polarities. Cells were initialized with random polarities.

**Supplementary Movie S11 –** Animated simulations of the model with pair-wise combinations of interactions; from top to bottom: VA+SPC, PA+SPC, VA+CRL, PA+CRL. Green dots: cell positions, green lines: elastic links between cells, magenta arrows: polarities. Cells were initialized with random polarities.

**Supplementary Movie S12 –** Animated simulations of the model combining velocity alignment, polarity alignment, and contact regulation of locomotion (VA+PA+CRL) and including cell-cell fragmentation beyond a critical length of 3, in a configuration mimicking the initial train formation. Cells at the back are attached to the slowly advancing tissue, while cells at the front are free to polarize. Green dots: cell positions, green lines: elastic links between cells, magenta arrows: polarities. Cells were initialized with random polarities.

**Supplementary Movie S13 –** Time-lapse sequence in differential Interference Contrast (DIC) mode of a cell cluster reaching the end of a 30 µm wide adhesive microstripe and entering a free open adhesive space. Upon exiting the microstripe, the cell cluster rapidly develops lamellipodia in the lateral directions away from their neighbours and finally breaks apart. The black arrow shows the direction of migration, while the light red zones indicate the microprinted adhesive areas.

**Supplementary Movie S14 –** Animated simulation of cell cluster reaching the end of a confined zone and entering a free open adhesive space.

**Supplementary Movie S15 –** Animated simulations of the model combining velocity alignment, polarity alignment, and contact regulation of locomotion (VA+PA+CRL). Green dots: cell positions, green lines: elastic links between cells, magenta arrows: polarities. Cells were initialized with random polarities. From top to bottom, systems with increasing width are shown, corresponding to n = 1,2,3,4 cells.

**Supplementary Movie S16 –** Time-lapse sequence in DIC mode of a cell train migrating on a microstripe of 15 µm wide with corners of 90°. The total duration time is 49 min. and the scale bar is 15 µm.

**Supplementary Movie S17 –** Animated simulations of the model combining velocity alignment, polarity alignment, and contact regulation of locomotion (VA+PA+CRL) and including cell-cell fragmentation in the weak (left) and strong (right) adhesion regimes. Geometries with a 90° angle are simulated. Green dots: cell positions, green lines: elastic links between cells, magenta arrows: polarities. Cells were initialized with random polarities.

**Supplementary Movie S18 –** Animated simulations of the model combining velocity alignment, polarity alignment, and contact regulation of locomotion (VA+PA+CRL), with parameters that capture the qualitative behavior of MDCK cell clusters. Top: microstripe (open boundary conditions). Bottom: microring (periodic boundary conditions). Green dots: cell positions, green lines: elastic links between cells, magenta arrows: polarities. Cells were initialized with random polarities.

**Supplementary Movie S19 –** Animated simulations of the repolarization behavior of model in a confined system with various interaction mechanisms. From top to bottom, the interaction mechanisms PA, VA, VA+CRL, VA+PA+CRL are shown. Green dots: cell positions, green lines: elastic links between cells, magenta arrows: polarities. Cells were initialized with random polarities.

**Supplementary Movie S20 –** Animated simulations of the model combining velocity alignment, polarity alignment, and contact regulation of locomotion (VA+PA+CRL) and including cell-cell fragmentation in the weak (left) and strong (right) adhesion regimes. Geometries with an open arena (top) and a blunt end (bottom) are simulated. Green dots: cell positions, green lines: elastic links between cells, magenta arrows: polarities. Cells were initialized with random polarities.

