## Supplementary Theory Note for "Geometry-driven migration efficiency of autonomous epithelial cell clusters"

In this Supplementary Theory Note, we provide additional details on the analytical and computational approaches used to model cell trains and clusters, as well as on our fitting strategy and comparison to the experimental data.

### 1 Model implementation

To identify the types of cell-cell interactions that capture the experimental phenomenology, we develop a minimal model of a polarizing active cell cluster. To be able to efficiently screen a wide range of possible cell-cell interactions, we seek a minimal model architecture that allows various couplings between cell polarities, velocities, and mechanical stresses to be easily implemented.

To this end, we model each cell as an active polarized particle, motivated by the observation that single keratocytes actively migrate with high persistence [1]. Cell-cell interactions naturally decompose into two classes: (i) positional cell-cell interactions, such as attraction and repulsion, which depend on relative cell positions; and (ii) orientational cell-cell interactions, which act on the cell polarities [2]. Since our cell cluster are confluent and exhibit few neighbour exchanges (see main text Figs. 1c, 2c and Supplementary Movies S1, S2, S5, S6), we assume a minimal implementation of positional cell-cell interactions through linear elastic couplings on an hexagonal lattice, which is a generic description as long as relative cell-cell displacement are small. With our model, we then seek to identify the orientational interactions that capture our experimental observations.

We therefore write the equations of motion of each cell $i$ in the cluster as

$$\dot{\mathbf{x}}_i = k \sum_{j \in \mathrm{NN}} \mathbf{f}(\mathbf{x}_j - \mathbf{x}_i) + \mathbf{p}_i(t) - \nabla V_{\text{confinement}}(\mathbf{x}_i). \tag{S1}$$

This makes the standard assumption that the migration dynamics is overdamped, hence that cell-substrate friction forces (left-hand term) balance cell-cell interactions and active migration forces (right-hand terms). The first right-hand term, with $\mathbf{f}(\mathbf{a}) = \mathbf{a}(1 - |\mathbf{a}|^{-1})$ gives the elastic force on cell $i$ due to all connected nearest-neighbour (NN) cells $j$, for elastic springs with spring constant $k$ and unit rest length. The second term represents active migration forces applied in the direction of polarity $p_i$. Finally, the third term represents geometrical confinement to the micropatterned lanes, which is implemented through the potential

$$V^{(y)}_{\text{confinement}} = ((y - \mathrm{sgn}(y) y_{\mathrm{w}})/y_{\mathrm{b}})^n \Theta(y_{\mathrm{w}} - y) \tag{S2}$$

where $\Theta$ is the Heaviside step function, and we set $y_{\mathrm{b}} = 0.25$, $n = 8$ and the width of the stripe proportional to the width of the cell cluster to $y_{\mathrm{w}} = (N_y - 1)/2$.

Although Eq. (S1) is a generic description of overdamped force balance for cell clusters, many types of orientational interactions have been proposed in the literature [2], and it was not straightforward to identify which of these would capture the observed experimental behaviour. We therefore decided to combine symmetry-based analytical theory with a computational screen of possible interaction types, to systematically test for their influence on migratory patterns. The dynamics of the cell polarity then read

$$\dot{\mathbf{p}}_i = \mathbf{p}_i(1 - |\mathbf{p}_i|^2) + \mathbf{F}^{\text{int}}_i + \sqrt{2D}\eta_i(t) \tag{S3}$$

Here, the first term enforces a polarized migration state (non-zero steady-state polarity), the second term subsumes a general class of possible interactions types that can contain couplings of cell positions, velocities and polarities, while the third term represents Gaussian white noise fluctuations of the polarity.

We then enumerate the possible orientational interaction terms $F^{\text{int}}$ that are allowed by symmetry up to first order (given the fact that polarity is a vector, and thus can be coupled only to vectorial quantities), and known from the cell migration literature. Specifically, we consider:

**Polarity alignment (PA)**: Here, each cell polarity tends to align to the average polarity of its neighbours ('Vicsek' alignment [3]).

$$\mathbf{F}_i^{\text{int}} = \beta_{\text{pol}} \left( \langle \mathbf{p}_j \rangle_{j \in \text{NN}} - \mathbf{p}_i \right) \tag{S4}$$

At the continuum level, this would correspond to a diffusional term applied to polarity [4].

**Velocity alignment (VA)**: Here, each single cell has a tendency to align its velocity vector to its polarity vector [5].

$$\mathbf{F}_i^{\text{int}} = \beta_{\text{vel}} \dot{\mathbf{x}}_i \tag{S5}$$

An alternative implementation is to align each polarity to the average velocity of the neighbouring cells. However, since cells are tightly bound in our model, cell velocities are influenced by the forces from the neighbors, and thus even single-cell velocity alignment results effectively in flocking. Thus, these two interaction types are equivalent within the framework of our model, and we therefore opt for the simple implementation of single-cell velocity alignment.

**Stress-polarity coupling (SPC)**: A prominent mechanism that causes anti-alignment between cells is stress-polarity coupling, which leads to cells polarizing away from forces. Based on symmetries, stress $\sigma_{ij}$ is a tensor and polarity $p_i$ can thus only be coupled to its divergence $\partial_j \sigma_{ij}$ (using the Einstein summation notation). Within our model, the only forces acting on each cell are the elastic couplings to its neighbours, so that this reads for discrete particles:

$$\mathbf{F}_i^{\text{int}} = -\lambda_{\text{SPC}} k \sum_{j \in \text{NN}} \mathbf{f}(\mathbf{x}_j - \mathbf{x}_i) \tag{S6}$$

where we explore $\lambda_{\text{SPC}} > 0$ since cells are typically considered to polarize in the opposite direction of an applied force [4,6]. Such stress-polarity coupling is furthermore typically considered to emerge in response to tensile stresses (pulling on the cell) rather than in response to pushing forces. We tested both implementations and did not find any significant differences, due to the fact that the stresses in the cluster are dominated by tensile stresses (see main text Fig. 4).

**Contact inhibition of locomotion (CIL)**: Another interaction mechanism that can cause anti-alignment between cells is contact inhibition of locomotion, where cells repolarize away from contacts with their neighbours [2,7]. Within our model, we implement this as a force that is directed away from the average of the unit vectors connecting cell $i$ with its neighbours:

$$\mathbf{F}_i^{\text{int}} = -\lambda_{\text{CIL}} \sum_{j \in \text{NN}} \frac{\mathbf{x}_j - \mathbf{x}_i}{|\mathbf{x}_j - \mathbf{x}_i|} \tag{S7}$$

In summary, with these four orientational cell-cell interactions, we consider the lowest-order couplings of the cell polarity dynamics to the four vectorial quantities in the problem: polarity itself (PA), velocity (VA), forces (SPC) or relative cell positions (CIL).

**Numerical simulation**: Numerically, the model is simulated using iterative Euler updates with time-interval $dt = 0.01$. Cell positions are initialized on a hexagonal grid and the corresponding connectivity matrix $c_{ij}$ is calculated which determines which cell $j$ is a nearest neighbour (NN) of cell $i$, used in Eqs. (S1), (S4), (S6), (S7). Polarities are initialized randomly with $x, y$ entries drawn from a Gaussian distribution, velocities are initialized to zero. In each simulation, we first run a pre-equilibration of 2500 time-steps to reach steady-state before statistics are recorded. We set $k = 10$ and $\sqrt{2D} = 0.05$ as standard values throughout, and vary the interaction parameters $\{\beta_{\text{vel}}, \beta_{\text{pol}}, \lambda_{\text{SPC}}, \lambda_{\text{CIL}}\}$.

**Dynamics in complex geometries**: We investigate the behaviour of cell clusters with various interaction rules in the presence of additional geometrical constraints in the lateral direction. Specifically, we consider the repolarization behaviour of cell clusters upon collision with the end of a microstripe and in the presence of a 90 degree turn. Both of these geometries are implemented by adding additional potentials with $x$-components in Eq. (S2). To quantify the efficiency of each interaction mode in terms of overall motion and speed of repolarization, we simulate a confined system and measure the average time taken to traverse the system $\tau_{\text{run}}$, and the average time taken to repolarize at the boundaries $\tau_{\text{rep}}$.

**Fragmentation simulations**: To investigate the conditions under which cell-cell contacts may rupture, we introduce a critical cell-cell contact extension $\ell_c$. If at any moment, the cell-cell distance of nearest neighbours exceeds this threshold, i.e. $|\mathbf{x}_j - \mathbf{x}_i| > \ell_c$, the contact fragments and these cells no longer interact through elastic forces or any other interaction.

**Implementation of cell-cell variability**: To investigate the cohesion of cell trains in the presence of variability of single-cell speeds, we include a minimal implementation of cell-to-cell variability in the single-cell polarity dynamics. Specifically, we scale the cell-autonomous polarity term in Eq. (S9) by an additional polarizability $\alpha_i$:

$$\dot{\mathbf{p}}_i = \alpha_i \mathbf{p}_i (1 - |\mathbf{p}_i|^2) + \mathbf{F}_i^{\text{int}} + \sqrt{2D}\eta_i(t) \tag{S8}$$

We then assume that the polarizability is normally distributed with standard deviation $\sigma_\alpha$:

$$\alpha_i \sim \mathcal{N}(\mu = 1, \sigma = \sigma_\alpha) \tag{S9}$$

### 2 Computational screen of possible interaction types

Based on our experimental observation that cluster speeds are constant with changing length, but decrease with width, we performed a parameter screen to identify the interaction types that can give rise to this combined set of observations. As shown in the main text (Fig. 2, Supplementary Movie S10), individual interaction types cannot capture these observations, as VA and PA alone predict geometry-independent speeds (i.e. where cells always flock at speeds similar to single cells, irrespective of geometry), while SPC and CIL predict speeds decreasing with both length and width (as trains with more than one cell in length develop bidirectional polarity patterns).

To explore our model, we first screened all pairwise combinations of these interactions. We initially reasoned that combining one interaction leading to flocking (PA, VA) with one leading to anti-flocking (SPC, CIL) could help to explain the data, for instance as each might react differently to the open boundary in the axial direction vs fixed boundary in the lateral direction.

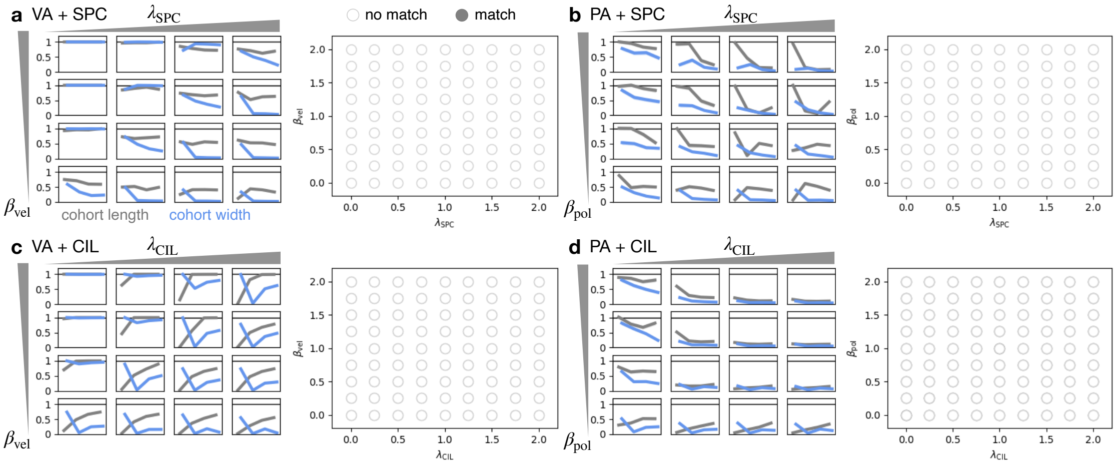


Figure S1: **Parameter sweeps of two-component combinations of cell-cell interactions. a.** Velocity alignment and stress-polarity coupling. **b.** Polarity alignment and stress-polarity coupling. **c.** Velocity alignment and contact inhibition of locomotion. **d.** Polarity alignment and contact inhibition of locomotion. *Left panels:* Dependence of the $x$-component speed, normalized by the speed of single cell, $v_x/v_x^{\text{single}}$ as a function of train length (blue data points, $N_x = \{1, 2, 4, 6, 8\}$) and train width (orange data points, $N_y = \{1, 2, 3, 4\}$). For each parameter, the values $\{0.25, 0.75, 1.5, 2\}$ are shown. *Right panels:* Phase diagram with a more detailed sweep. Open symbols correspond to parameter combinations where the experimental phenomenology is not captured (defined as speed being constant as a function of $N_x$, and decreasing as a function of $N_y$).

However, among pairwise combinations of interactions, no combination could capture the experimental observations (Fig. S1, Supplementary Movie S11). For both VA+SPC and PA+SPC, we find that for SPC couplings large enough to cause a decrease of speed with width, speeds also decrease with length (Fig. S1a,b). This is because SPC has no coupling to geometry in the sense that it causes the same amount of anti-alignment in both the axial and lateral directions. Accordingly, for one-dimensional cell trains with SPC, we find misaligned polarities, in disagreement with experiment.

For VA + CIL, we find a non-monotonic dependence of speed with width (Fig. S1c). This is because here, only the outermost lateral layer experiences outward polarization in the $y$-direction due to CIL, while the inner layers can completely align with the axial direction of motion. Thus, the $N_y = 2$ state is completely non-motile for many parameter combinations.

For PA + CIL, we find again that for CIL coupling strengths large enough to cause a decrease of speed with width, speeds also decrease with length (Fig. S1d). This is because for the one-dimensional cell trains, the first and final cells are affected by CIL and the alignment of polarities causes the center cells to redirect their polarities in non-axial directions to reduce polarity gradients, causing less polarization in the axial direction of motion.

Our observations on the pairwise interactions of VA and PA with CIL already indicate that a combination of these three interactions has the potential to capture the data: VA + CIL can capture constant speed with length, with a non-monotonic width-dependence. Polarity alignment can cause inward-propagation of the CIL-induced outward pointing polarities at the lateral boundaries, which can lead to a decreasing trend of speed with width. Indeed, we find that the experimental observations are captured in a model including VA + PA + CIL in a parameter regime where the magnitudes of velocity alignment and contact inhibition of locomotion are approximately balanced $\beta_{\text{vel}} \approx \lambda_{\text{CIL}}$ and both are smaller than the magnitude of polarity alignment $\beta_{\text{pol}}$ (Fig. S2). In contrast, we find that the combinations of VA + PA + SPC and of VA + SPC + CIL do not capture the experiments (Fig. S3, S4). Finally, we examine the parameter regimes with a qualitative match and search for parameters that can quantitatively capture the decay of speeds with width to within error, and find quantitative matches for parameter regions corresponding to below the diagonal in Fig. S2b, i.e. where $\beta_{\text{vel}} < \lambda_{\text{CIL}}$. We find the best match for $\beta_{\text{pol}} = 1.5$, $\beta_{\text{vel}} = 0.15$, $\lambda_{\text{CIL}} = 0.5$, which is used in Fig. 2 and subsequently - with no further fitting - for the prediction of the monolayer stresses.

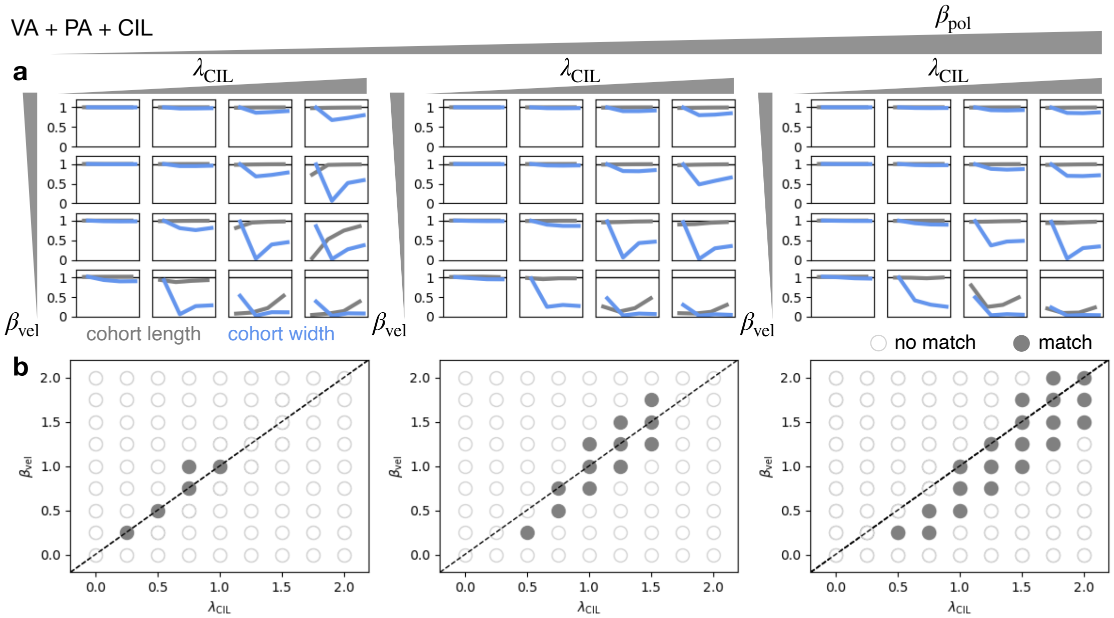


Figure S2: **Parameter sweeps of the model with combined velocity alignment, polarity alignment and contact inhibition of locomotion.** The polarity alignment strength increases from left to right with values $\beta_{\text{pol}} = \{1, 1.5, 2\}$ **a.** Dependence of the $x$-component speed, normalized by the speed of single cell, $v_x/v_x^{\text{single}}$ as a function of train length (blue data points, $N_x = \{1, 2, 4, 6, 8\}$) and train width (orange data points, $N_y = \{1, 2, 3, 4\}$). For each parameter, the values $\{0.25, 0.75, 1.5, 2\}$ are shown. **b.** Phase diagram with a more detailed sweep. Open symbols correspond to parameter combinations where the experimental phenomenology is not captured (defined as speed being constant as a function of $N_x$, and decreasing as a function of $N_y$).

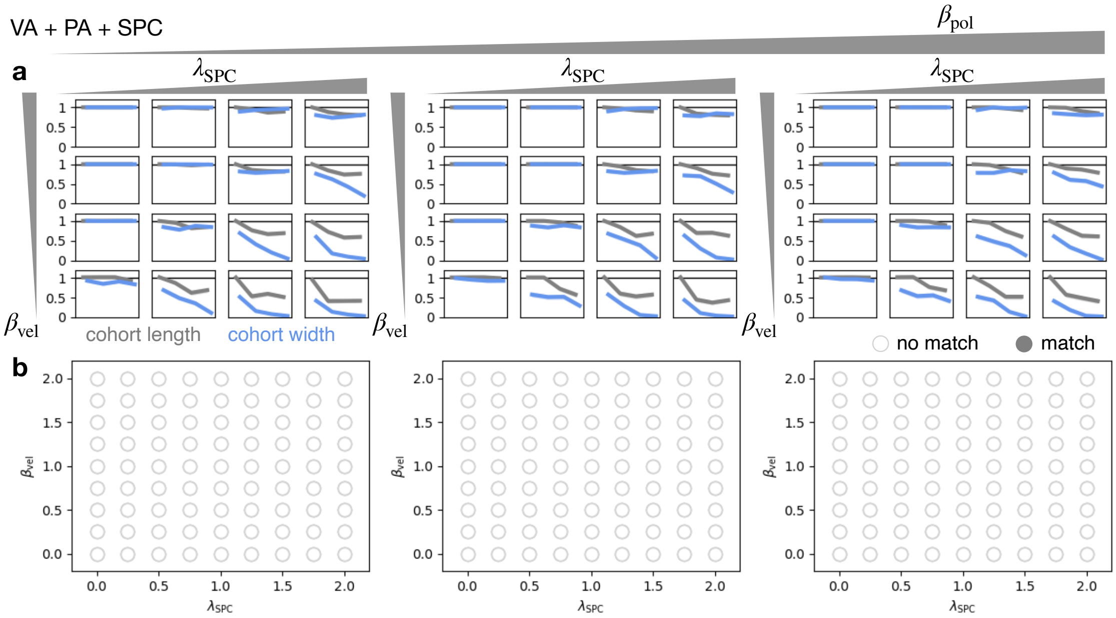


Figure S3: **Parameter sweeps of the model with combined velocity alignment, polarity alignment and stress-polarity coupling.** The polarity alignment strength increases from left to right with values $\beta_{pol} = \{0.5, 1.5, 2\}$ **a.** Dependence of the $x$-component speed, normalized by the speed of single cell, $v_x/v_x^{single}$ as a function of train length (blue data points, $N_x = \{1, 2, 4, 6, 8\}$) and train width (orange data points, $N_y = \{1, 2, 3, 4\}$). For each parameter, the values $\{0.25, 0.75, 1.5, 2\}$ are shown. **b.** Phase diagram with a more detailed sweep. Open symbols correspond to parameter combinations where the experimental phenomenology is not captured (defined as speed being constant as a function of $N_x$, and decreasing as a function of $N_y$).

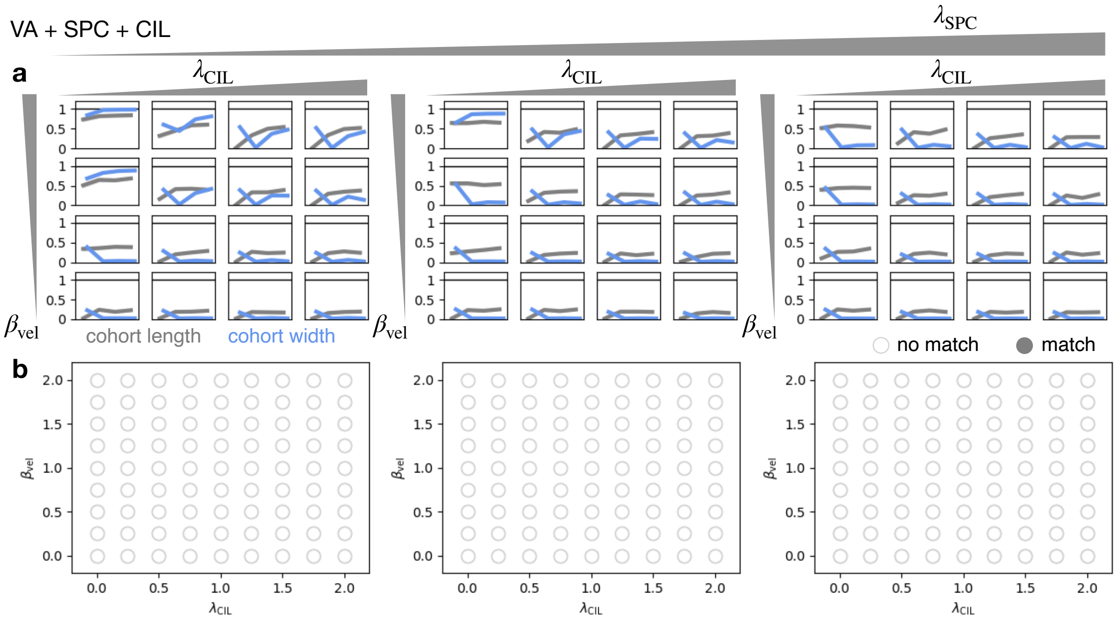


Figure S4: **Parameter sweeps of the model with combined velocity alignment, contact inhibition of locomotion and stress-polarity coupling.** The polarity alignment strength increases from left to right with values $\beta_{pol} = \{0.5, 1.5, 2\}$ **a.** Dependence of the $x$-component speed, normalized by the speed of single cell, $v_x/v_x^{single}$ as a function of train length (blue data points, $N_x = \{1, 2, 4, 6, 8\}$) and train width (orange data points, $N_y = \{1, 2, 3, 4\}$). For each parameter, the values $\{0.25, 0.75, 1.5, 2\}$ are shown. **b.** Phase diagram with a more detailed sweep. Open symbols correspond to parameter combinations where the experimental phenomenology is not captured (defined as speed being constant as a function of $N_x$, and decreasing as a function of $N_y$).

### 3 Simulations of cluster fragmentation and cell-to-cell variability

To check that the constant cell speed of cellular trains was not an artifact due to cells of different intrinsic speed detaching from each other, we included the possibility of junctional breakage in our model (when cell-cell distance exceeds a critical threshold $\ell_c$) as well as intrinsic variability in cellular migration forces (based on single-cell migration speed variability, Fig. S5a,b). For this, we first infer the amplitude of the variability in single cell speeds in the experiments. We find significant cell-to-cell variability in single-cell speeds, with an average standard deviation of $\sim 48\%$ of the average speed ($8.45 \pm 4.05$ $\mu$m/min, Fig. S5a,b). As a minimal implementation of this variability in the model, we introduce a prefactor $\alpha$ in the polarity force of each cell, which is randomly distributed across single cells (see section 1). This leads to a distribution of single cell speeds, with an amplitude fitted to that observed experimentally (Fig. S5c). In the regime of weak adhesion ($\ell_c = 1.2\ell$), such speed variability leads to rapid fragmentation of the cluster (Fig. S5d). However, using the experimentally inferred value of the critical length ($\ell_c = 3\ell$, based on the fragmentation in the initial fingering process; Extended Data Fig. 1) we predict that the clusters do not fragment (Fig. S5e,f), which is consistent with our experimental observation that cell trains very rarely fragment after initial train formation. Using the experimentally inferred values of critical length and cell-to-cell variability in speed, we then simulate trains of varying length, and find that their average speed is constant with length, albeit with a gradually decreasing width of the train speed distributions, due to increased averaging from cell-cell interactions. Taken together, this shows that based on the experimentally inferred speed variability and adhesion strength, we do not expect train fragmentation, and predict constant speed with length, as observed experimentally.

Furthermore, we also used this new model to make predictions for various other experimental geometrical confinements. Firstly, for the experiments where a confined cluster migrates into an open arena, we find experimentally that the cell trains de-compress, due to the outwards-directed cell polarities. In the weak adhesion regime, our model predicts that this would lead to a fragmentation of the cell clusters into individual cells or smaller clusters (Fig. S6a, Supplementary Movie S20). In contrast, in the strong adhesion regime parametrized above, the cluster remains cohesive, but develops outward polarities leading to tumbling of the cluster (Fig. S6b). This latter prediction is consistent with our experimental observations (Extended Data Fig. 9). Secondly, similar simulations for systems with a blunt end of the microstripe predict rupturing of the cluster upon turning around, as the back cells repolarize faster, generating velocity differences in the cluster (Fig. S6c, Supplementary Movie S20). This does not occur in the strong adhesion regime, and is also not observed experimentally (Fig. S6d). Finally, we test the behaviour of cell clusters navigating around 90 degree corners. In the regime of weak adhesion, cell clusters indeed break apart when migrating into a 90 degree turn, due to the large velocity gradients induced by the geometry (Fig. S6e, Supplementary Movie S17). In contrast, for strong adhesion, cell clusters are predicted to remain cohesive and rapidly reorient, consistent with our experimental observations (Fig. S6f).

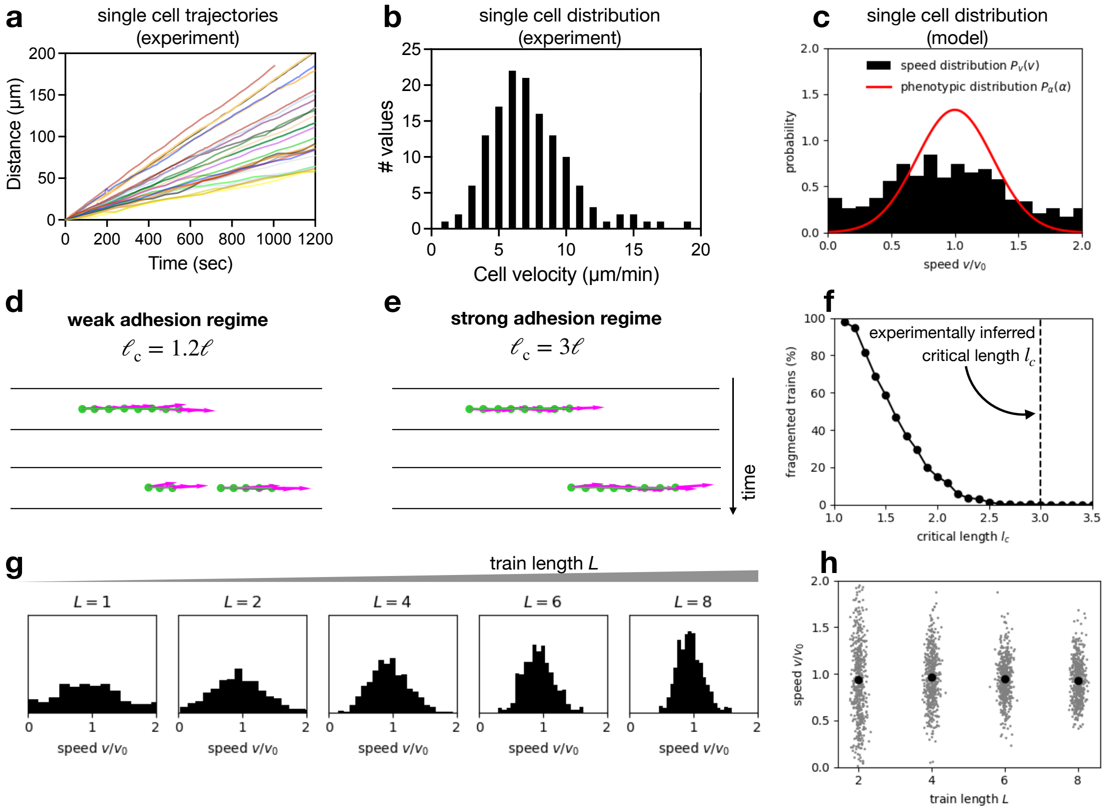


Figure S5: **Predictions of the fragmentation model with experimentally inferred cell-to-cell speed variability. a.** Experimental single cell trajectories, showing low stochasticity in a single trajectory, but large variability across cells. **b.** Distribution of single-cell speeds, providing an estimate of cell-to-cell variability in speeds. **c.** Using the phenotypic distribution of the polarity prefactor $\alpha$ (line) in the model, we obtain a broad distribution of single-cell speeds (histogram). **d,e.** Time-series of simulated cell train migration, including cell-to-cell variability, in the weak and strong adhesion regimes, respectively. This shows fragmentation in the weak regime and cohesive migration in the strong regime. **f.** Percentage of fragmented trains in simulations with varying critical junction length lc. Dashed line indicates the experimentally inferred critical junction length from the initial fingering process. **g.** Speed distributions of cell trains of varying length L in the strong adhesion regime. **h.** Train speed as a function of length, showing constant mean speed despite the presence of cell-to-cell variability in single-cell speeds.

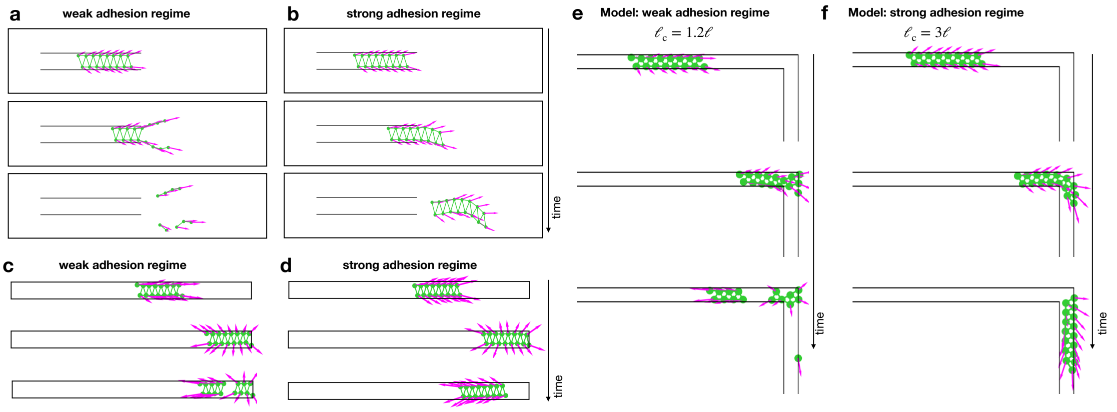


Figure S6: **Predictions of the fragmentation model in various conditions. a,b.** Simulations of cell clusters exiting into an open arena in the weak and strong adhesion limit, showing fragmentation of the cluster in the weak adhesion regime, while in the strong adhesion regime, clusters remain cohesive. **c,d.** Simulations of cell clusters migrating into a blunt end in the weak and strong adhesion regime, showing fragmentation in the weak adhesion regime. **e,f.** Simulation of cell train migrating around corners of 90 degrees. In the weak adhesion regime, clusters fragment upon colliding with the corner. In the strong adhesion regime, cell clusters remain cohesive and efficiently repolarize around the corner.
