## Supplementary Information for "Geometry-driven migration efficiency of autonomous epithelial cell clusters"

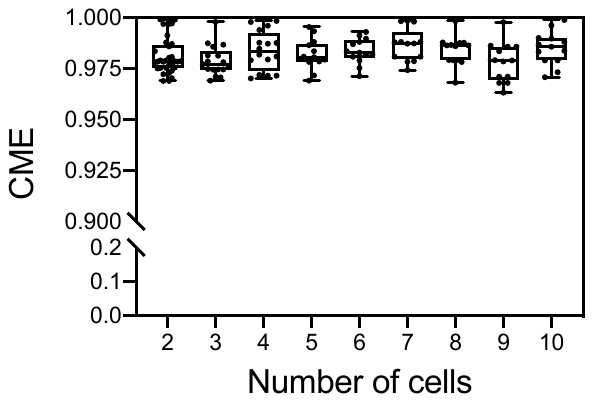

**Supplementary Figure 1 – Persistence of cell trains.** The coefficient of movement efficiency (CME), corresponding to the ratio of the total cell displacement (TCD) to the cell trajectory length (CTL) was determined for cell trains of following lengths: L=2 (n=32), L=3 (n=17), L=4 (n=16), L=5 (n=13), L=6 (n=13), L=7 (n=13), L=8 (n=13), L=9 (n=13) and L=10 (n=13). A minimum of 3 replicates was used for each condition.

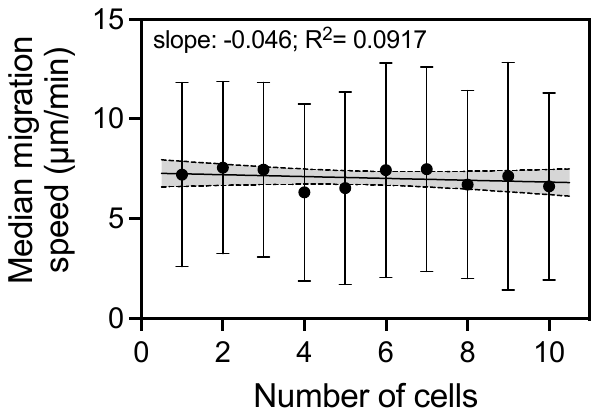

**Supplementary Figure 2 – Median migration speed versus the number of cells.** The linear regression (black line) of the median cell train speed of length 1≤L≤10 shows a slope of -0.046±0.07, indicating no trend between the cell train migration speed and the train length. Data are shown as median ± interquartile range and dashed lines represent the 95% confidence bands of the best-fit line (R^2^=0.0917).

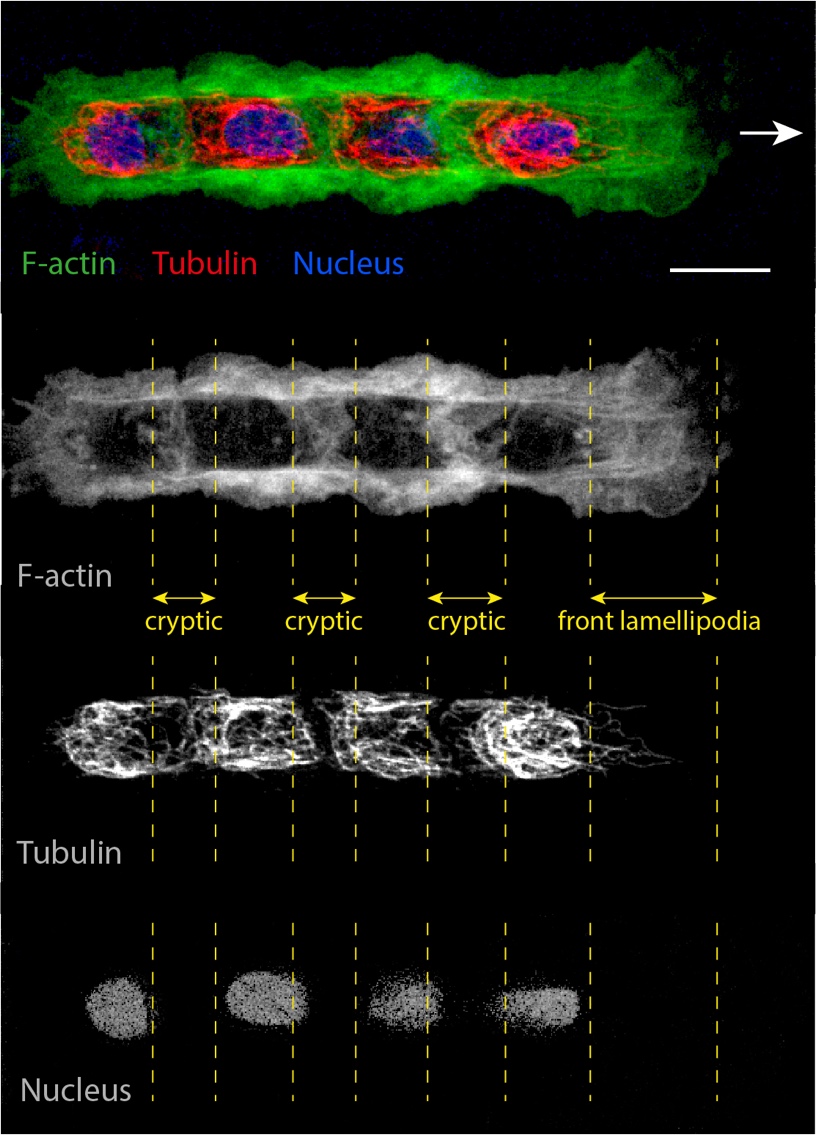

**Supplementary Figure 3 – Spatial distribution of the microtubule network in cell trains.** Epifluorescent image of a train of L=4 cells migrating on a fibronectin microstripe of 15 µm wide and stained for tubulin. Keratocytes were labelled for the actin network (green), microtubules (red) and the nucleus (blue). The direction of motion is depicted by the white arrow. and the orientation of the Golgi complex relative to the nucleus is indicated by a yellow arrow. The vertical yellow dashed lines represent the positions of the cryptic and front lamellipodia. The scale bar is 15 µm.

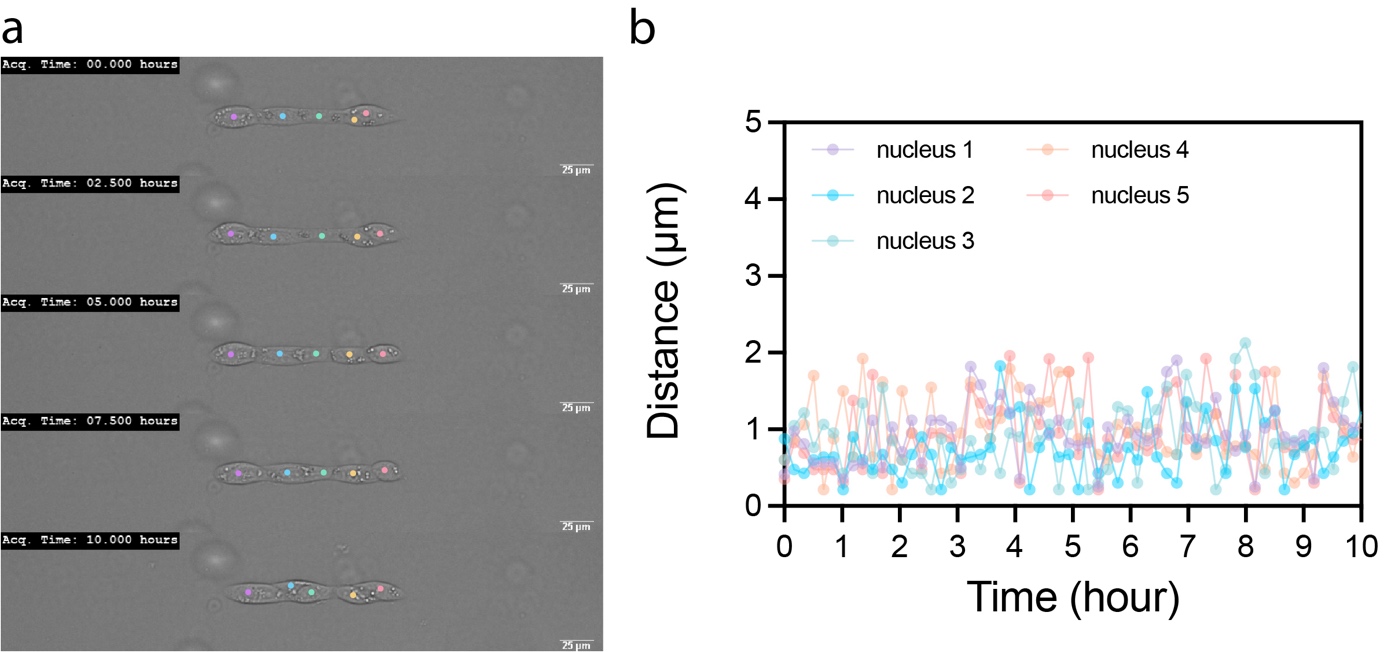

**Supplementary Figure 4 – Behvaiour of autonomous MDCK cell trains.** (a) Time-lapse sequence of a MDCK train of L=5 cells on a microstripe of 15 µm wide for 10 hours. The center of mass of each nucleus is represented by a color spot. (b) The temporal evolution of the center of mass of each nucleus showed that they fluctuated by only a few microns per hour, indicating the MDCK cell train remained stationary.
